# Two therapeutic CRISPR/Cas9 gene editing approaches revert FTD/ALS cellular pathology caused by a *C9orf72* repeat expansion mutation in patient derived cells

**DOI:** 10.1101/2022.05.21.492887

**Authors:** M Sckaff, K Gill, A Sachdev, AM Birk, O Aladesuyi Arogundade, HL Watry, KC Keough, Y-C Tsai, J Ziegle, BR Conklin, CD Clelland

**Affiliations:** Weill Institute for Neurosciences, University of California San Francisco, San Francisco, California 94158, USA; Gladstone Institutes, San Francisco, California, 94158, USA; Pacific Biosciences, Menlo Park, California 94025, USA; Departments of Medicine, Ophthalmology, and Pharmacology, University of California San Francisco, San Francisco, California 94143, USA; Memory & Aging Center, Department of Neurology, University of California San Francisco, San Francisco, California 94158, USA

## Abstract

CRISPR gene editing holds promise to cure or arrest genetic disease, if we can find and implement curative edits reliably, safely and effectively. Expansion of a hexanucleotide repeat in *C9orf72* is the leading known genetic cause of frontotemporal dementia (FTD) and amyotrophic lateral sclerosis (ALS). We evaluated three approaches to editing the mutant *C9orf72* gene for their ability to correct pathology in neurons derived from patient iPSCs: excision of the repeat region, excision of the mutant allele, and excision of regulatory region exon 1A. All three approaches normalized RNA abnormalities and TDP-43 pathology, but only repeat excision and mutant allele excision completely eliminated pathologic dipeptide repeats. Our work sheds light on the complex regulation of the *C9orf72* gene and suggests that because of sense and anti-sense transcription, silencing a single regulatory region may not reverse all pathology. Our work also provides a roadmap for evaluating CRISPR gene correction using patient iPSCs.

## Introduction

Age-related neurodegenerative diseases, including dementias and motor neuron diseases, are leading contributors to death, disability and health care expenditure worldwide^1–6^. Heterozygous expansion of a GGGGCC repeat in the *C9orf72* gene is the most frequent known genetic cause of both frontotemporal dementia (FTD) and amyotrophic lateral sclerosis (ALS)^7–9^ (C9FTD/ALS). Targeting the mutant *C9orf72* gene itself is the most parsimonious and potentially the most powerful therapeutic intervention. While antisense oligonucleotide (ASO) therapy showed promise in pre-clinical studies^10–14^, clinical futility of a phase I ASO trial in C9-ALS patients^15^ demonstrates the need for other approaches. Gene editing offers the advantage that a single intervention could be curative^16^.

C9FTD/ALS pathology is thought to result from the generation of toxic products derived from expression of the *C9orf72* repeat expansion itself. The resulting RNA harbors a large repeat expansion that produces toxic dipeptide repeat proteins (DPRs) through repeat-associated non-canonical (RAN) translation^17–26^, and may disrupt RNA processing by sequestering RNA-binding proteins^22, 27–30^. Haploinsufficiency has been proposed as an additional or alternative mechanism of disease^31–34^, but it is unlikely to be the major contributor to C9FTD/ALS. The most compelling evidence against this hypothesis is that large-scale population sequencing^35^ and clinical sequencing suggest that *C9orf72* heterozygous loss-of-function mutations do not cause C9FTD/ALS^36^. Secondly, knock-out mouse models have an autoimmune phenotype but lack neurologic disease^37–40^. On the other hand, loss of *C9orf72* function may exacerbate the toxic gain-of-function^39, 41^ caused by the repeat expansion. Since pathology appears to result mainly from the mutant allele, and harboring a single, non-expanded copy of the *C9orf72* gene is well tolerated, we elicited to reverse pathology by editing the expanded allele.

What is the editing strategy with the best therapeutic potential? The simplest strategy might be to shorten the repeat expansion itself^42–45^. But this approach carries the risk of off-target editing at more than 2500 sites throughout the genome that have homology to the GGGGCC motif^46^, or of cellular death from widespread DNA damage. Other approaches disrupt regulatory regions on both the normal and diseased allele^42, 45^, which could be deleterious as homozygous knockout causes early lethality in mice^37–39^. Finally, editing strategies that utilize homology directed repair^44^ are inefficient in post-mitotic cells^47^ such as neurons, and are therefore not suitable for therapeutic applications.

We designed our approaches to avoid these pitfalls, taking advantage of the CRISPR/Cas9 system as a highly specific genome editing tool and of newly engineered Cas9 variants capable of distinguishing alleles differing by even a single base pair^48–55^. Here we report our success at reverting hallmarks of disease, mRNA abnormalities and loss of nuclear TDP-43, in human motor neurons derived from diseased iPSCs with three strategies: (1) bi-allelic excision of the repeat region, (2) allele-specific excision of the mutant allele leaving the normal allele intact and (3) excision of a regulatory region (exon 1A) that controls expression of the mutant allele’s sense-strand. Only the two strategies that removed the repeat expansion itself also eliminated dipeptide repeat expression. We propose single-molecule sequencing as a gold standard for determining editing outcome of this repetitive region. Our work also revealed aspects of gene expression and regulation at the *C9orf72* locus that should inform future gene editing approaches.

## Results

### Single-molecule sequencing distinguishes repeat expansion lengths in patient-derived iPSC lines

Currently, long-range PCR^56^ and Southern blot^57^ are used to clinically diagnose repeat expansion mutations in the *C9orf72* gene, but these approaches lack in precision, especially for large expansion. Sizing the repeat expansion above ∼100 repeats is not possible using traditional sequencing techniques that require amplification because amplification fails across GC-rich repetitive DNA regions. Patients can have *C9orf72* repeats into the thousands! We turned instead to single-molecule sequencing, which has been demonstrated to traverse the expanded repeats of *C9orf72* in plasmid^58^ and human tissue^59, 60^. We collected patient iPSC lines from previously published or publicly available sources^31, 42, 61, 62^ and developed PacBio single-molecule sequencing of DNA from these lines to size their repeat expansions and identify surrounding SNPs (**Extended Data Fig. 1**). Using Cas9 to generate double-stranded breaks (DSBs), adapter ligation to capture the cut genomic region of interest, and exonuclease digestion to eliminate DNA fragments without adapter, we were able to enrich for a 3.6-10kb genomic region centering on the repeat expansion without amplification (**Extended Data Fig. 1a**). Sequencing these single, circular DNA fragments gave us a more precise count of the number of repeats present in each cell line (**Extended Data Fig. 1b**) than Southern blots did (**Extended Data Fig. 1c**). In addition, single molecule sequencing was more sensitive, requiring an input of only 3 µg DNA versus 20 µg for Southern blot.

From these data, we chose a patient cell line with ∼200 repeats in one allele and 2 repeats in the other (Patient 3, **Extended Data Fig. 1b,c,j,k)**. This line also features an advantageous SNP in the splice acceptor of exon 2, which we later exploited for measuring the RNAs produced by each allele. From here on out, we refer to this line as C9-unedited. A cell line from a non-diseased patient with 2 repeats on one allele and 10 on the other serves as our control and is referred to as WT-control.

### Short excisions near the repeat expansion are less efficient than a 22 kb excision of the mutant allele

The *C9orf72* mutation lies in the non-coding 5’UTR between two alternative start sites, exon 1A and exon 1B^7, 8^ (**Fig. 1a**). We hypothesized that gene editing strategies that can remove or silence the repeat expansion would be curative at the cellular level. We compared three editing approaches to correcting the *C9orf72* mutation in our C9 and WT cell lines (**Fig. 1b,c**). Each of these approaches capitalizes on the ability of Cas9 to induce double-stranded breaks (cuts) in DNA, which aligns with the most-developed Cas9 technology currently employed in clinical trials^63–65^. We used CRISPOR^46, 66^ or AlleleAnalyzer^67^ to design gRNAs (**Supplementary Table 1**) with the fewest computationally predicted overall off-targets (**Supplementary Table 2**), including no predicted off-target matches to the exact gRNA sequence and no predicted off-target within the first 2 bases of the PAM.

**Figure 1.**
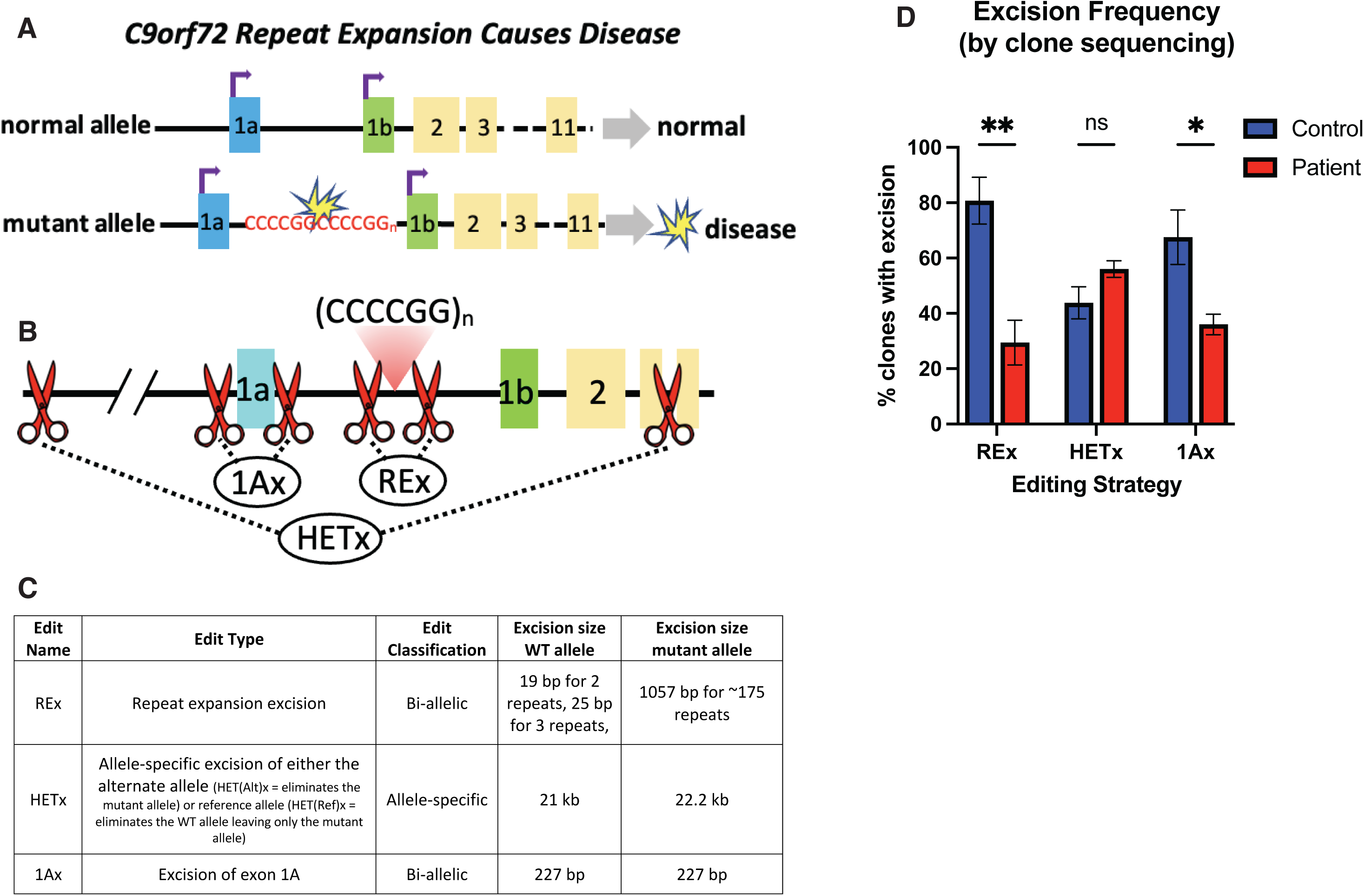
Editing efficiencies of three therapeutic CRISPR editing approaches of editing the repeat region of the *C9orf72* locus in non-disease control and patient iPSC lines. (A) The repeat region of the *C9orf72* gene lies in the 5’ UTR, between alternative start sites that give rise to non-coding exon 1a (blue) and exon 1b (green). Translation starts in exon 2 (yellow). Only the allele with the repeat expansion causes disease. (B) Three CRISPR gene editing approaches to correct the *C9orf72* mutation. REx: bi-allelic removal of the repeat expansion region. HETx: large allele-specific excision spanning the repeat expansion and both transcriptional start sites. This approach leaves the normal allele intact. 1Ax: Bi-allelic excision of exon 1A, which controls expression of a sense mRNA containing the repeat expansion. (C) Editing approaches and expected excision sizes for the mutant and WT alleles. (D) Editing efficiencies were determined by PCR or single-molecule sequencing across the excision site. Each experiment contained 3 biologic replicates of either 48 hand-picked or 96 single-cell sorted clones. Bi-allelic excision of the repeat expansion (REx) or of intron 1A (1Ax) was significantly less efficient in patient than in control lines, whereas the efficiency of the large allele-specific gene excision (HETx) was similar in both backgrounds (2-way ANOVA F (2, 11) = 9.115, p<0.001; *p<0.05, **p<0.01 using Sidaks multiple comparison post-hoc test). Error bars = SEM.

Our first approach excised the repeat expansion region (REx, **Fig. 1c, Extended Data Fig. 3**). Given numerous predicted off-targets throughout the genome, it is not safe to cut within the repeat region itself; instead we made cuts just 5’ and 3’ to the repeat region (**Fig. 1b,** REx). This area is highly conserved and does not offer allele-specific gRNA binding sites. Therefore this excision was expected to be bi-allelic. However, by chance it occurred only on the mutant allele in our patient cell line (**Extended Data Fig. 3**), leaving intact the native two repeats on the WT allele. Our second approach was to excise the mutant allele, leaving the normal allele intact (**Fig. 1b,** HETx). Newer versions of Cas9 can distinguish between alleles that differ by a single nucleotide^48^, which allowed us to use SNPs in cis with the mutation to specifically excise the mutant allele. We used AlleleAnalyzer^67^, an open source bioinformatics tool, to design allele-specific gRNA pairs based on common heterozygous polymorphisms found in a reference dataset of over 2500 human genomes from around the world^68^. We chose a pair of gRNAs (**Supplementary Table 1**) that would excise a large segment of the *C9orf72* locus, cover the maximum number of individuals in the representative global cohort, and have the lowest number of predicted off-targets. Using this pair, we excised 22.2 kb of the mutant allele (HET(Mut)x) starting 12.3 kb upstream of exon 1A and stretching all the way through exon 3 (**Fig. 1b,c,** HETx; **Extended Data Fig. 4**). In addition, we made the equivalent 21 kb excision on the WT allele, leaving the mutant allele intact (HET(WT)x) (**Extended Data Fig. 7**). Our third approach was to leave the mutation in the DNA but silence its expression by excising exon 1A, which includes a transcriptional start site and controls the expression of the *C9orf72* sense-transcript harboring the mutation (**Fig. 1b,c,** 1Ax, **Extended Data Fig. 5**). We also made the REx, HETx and 1Ax excisions in a WT-control line to examine the effects of each approach on the normal cellular expression of the *C9orf72* gene (**Extended Data Figs. 8-10**). As additional controls, we also created homozygous knock-outs of the gene in our patient and WT lines, using bi-allelic excisions starting upstream of exon 1A and ending in exon 3 or exon 2, respectively (**Extended Data Figs. 6,11**).

We measured editing efficiency (**Fig. 1d**) by PCR or single-molecule sequencing across the edited locus in 3 independent experiments per edit. Each experiment derived from 48 hand-picked or 96 single-cell sorted clones. The efficiency of all editing was between 21 and 92% in iPSCs. We found that editing near the repeat region (REx, 1Ax) was significantly less efficient in the patient lines containing a large repeat expansion than in the WT-control line with fewer than 10 repeats. One hypothesis is that methylation of the repeat region and promoter^69–71^ in patient lines reduces access to the loci for Cas enzymes and therefore lowers editing efficiency. Interestingly, the efficiency of the large (21 or 22.2 kb) excision was surprisingly high (30-59%) and did not differ between patient and WT lines. Further improvements to the efficiency of the various edits might yet be possible, as the gRNAs used here were selected strictly for their predicted on- and off-target rates, and have not been optimized experimentally for efficiency.

### Single-molecule sequencing affords better resolution of editing outcomes after dual gRNA excisions than Sanger sequencing

Single-molecule sequencing was critical to determining the outcomes of edits targeting the expanded allele (**Extended Data Fig. 3**). Using Sanger sequencing, it is not possible to determine whether an edit removed the repeat region from both the expanded and WT alleles or just from the WT allele, since the mutant allele fails amplification, and hence detection. Because of its low sensitivity, Southern blot may not reveal the impurity of a cell line harboring a mix of mutant and edited clones. With high sensitivity and ability to read through long repeat expansions, we find single-molecule sequencing to be superior to current ways of determining editing outcome at the *C9orf72* locus, and recommend it become the new gold standard for verifying lines with *C9orf72* edits. In addition, single-molecule sequencing is more amenable to SNP detection and has the ability to phase SNPs (**Extended Data Fig. 3d**). It can also detect very small or very large excisions, inversions and introductions of single nucleotide variants, all of which are hard to resolve by Sanger sequencing. For example, Sanger sequencing could not resolve whether a 1 bp shift at the 5’ cut site of WT 1Ax reflected a mixed population of two clonal lines that differed by a single nucleotide or a single clonal line with a one-nucleotide difference between its alleles. Using nearby SNPs to specify each allele, single-molecule sequencing quickly showed our WT 1Ax line had a 1-nucleotide deletion on one allele (**Extended Data Fig. 10d**).

### The *C9orf72* 1A-transcriptional start site is more active, and the 1B-transcriptional start site less active, in the mutant than the WT allele

We next evaluated the effect of each of the edits on *C9orf72* RNA expression. The *C9orf72* locus is known to produce at least three sense mRNAs: variant 1 (exon 1A-short through exon 5), variant 2 (exon 1B-exon 11) and variant 3 (exon 1A-long through exon 11)7. Using ddPCR probes spanning either the exon 1A-exon 2 or exon 1B-exon 2 splice junctions, we quantified the two major splice forms of *C9orf72*, variant 3 and variant 2, which we refer to as 1A-transcript and 1B-transcript from here on (**Fig. 2a,b; Supplementary Table 3**). We were not able to detect variant 1 in our lines, consistent with its low to undetectable expression in human tissue^72, 73^. We also quantified total *C9orf72* mRNA using a probe targeting the exon 2-exon 3 junction.

**Figure 2.**
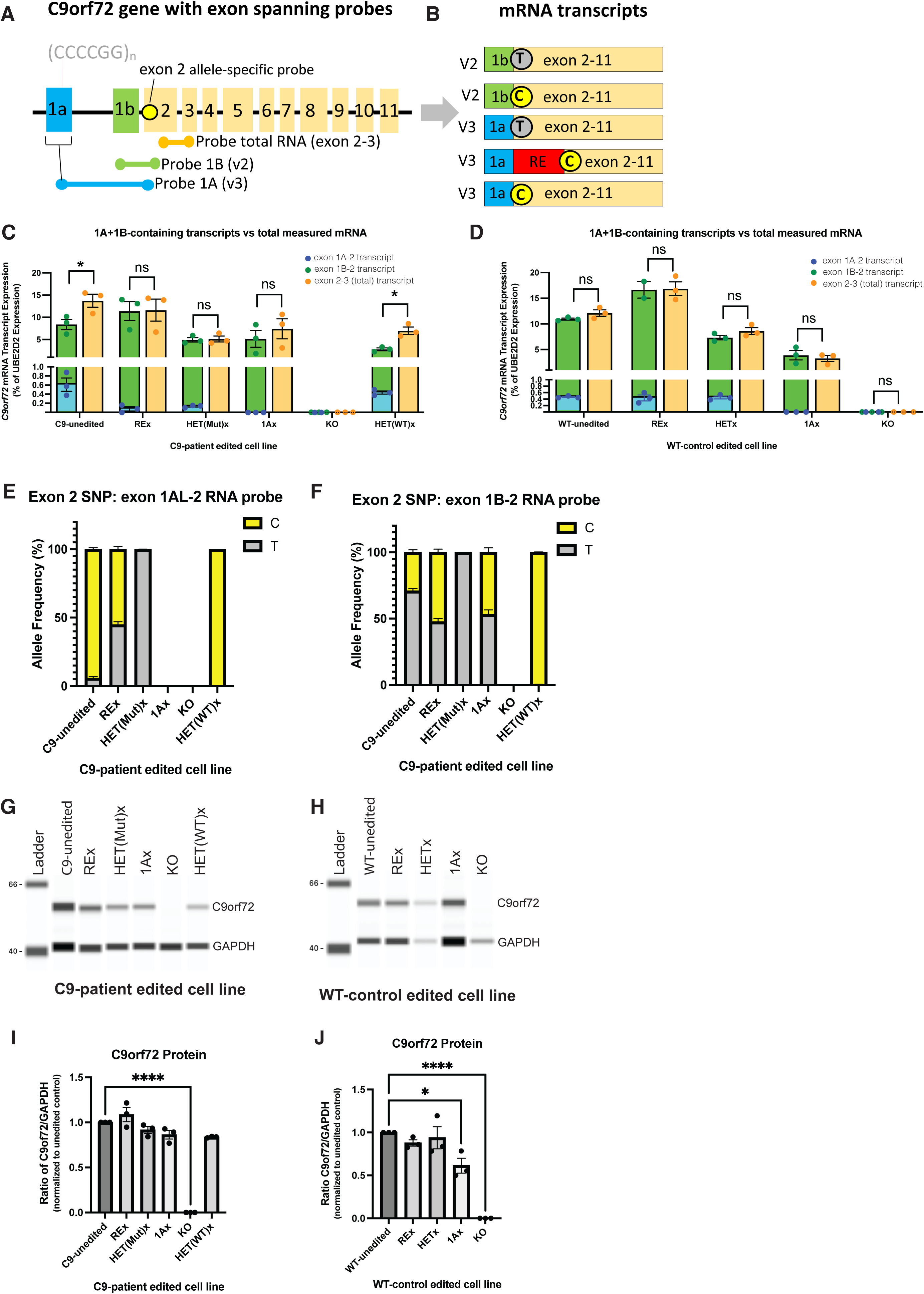
*C9orf72* gene expression after gene editing in 2-week-old hNIL neurons. (A,B) Location of ddPCR probes (A) and schematic of the possible allele-specific transcripts (B). ddPCR probes are designed to distinguish mRNA transcripts starting at exon 1A (V3) versus exon 1B (V2), or to detect total *C9orf72* mRNAs (probe spanning exons 2-3). An additional ddPCR probe spans a SNP in exon 2 that we used to distinguish transcripts from the wild-type allele (carrying a T) versus the mutant allele (carrying a C). Exon 1a normally splices onto exon 2, but the repeat expansion disrupts this splicing event. Exon 1A transcripts from the C-allele are only detected if correctly spliced, due to amplification failure of the repeat region. (C,D) ddPCR quantification of exon 1A-containing RNA (blue), exon 1B-containing RNA (green) and total *C9orf72* RNA (orange) in isogenic lines from a C9-patient (C) or WT-control (D), normalized to expression of the *UBE2D2* housekeeping gene. In all cell lines, the majority of *C9orf72* mRNAs start at exon 1B, and mRNAs starting at exon 1A are 20 to 100 times less abundant than those starting at exon 1B. In lines harboring a repeat expansion (C9-unedited, HET(WT)x), the sum of exon-1A- + exon-1B- containing transcripts was significantly smaller than the amount of total transcripts (containing exons 2-3) (paired t-test corrected for multiple tests, FDR<5%, *=p<0.01). We hypothesize that this gap corresponds to improperly spliced 1A RNA from the mutant allele, since the presence of a repeat expansion would disrupt exon 1A-exon 2 primer amplification as well as probe binding. (E, F) Proportion of 1A (E) and 1B (F) transcripts coming from the C versus T allele in unedited and edited C9 lines, as determined by ddPCR. The majority of 1A RNA came from the mutant allele in the C9 unedited line and this was normalized by excision of the repeat expansion (E, REx). Reciprocally, 1B RNA predominantly arose from the WT allele in the unedited patient line; this imbalance was corrected by repeat expansion excision (REx) or excision of 1A (1Ax). As expected, heterozygous excision of either allele resulted in expression off of the other (E,F: HET(Mut)x and HET(WT)x), bi-allelic excision of exon 1A (1Ax) abolished expression 1A but not 1B transcripts, and bi-allelic gene knock-out (KO) abolished all transcripts). (G-J) Quantification of C9orf72 protein expression by Simple Western (WES) relative to GAPDH in unedited and edited C9 (G, I) and WT (H, J) lines. C9orf72 protein levels remained constant in both backgrounds after edits, except after biallelic 1A excision in WT, or after homozygous gene KO in both C9 and WT lines. (1-way ANOVA: C9: F(5,12)=94.81, p<0.0001; WT: F(4,10)=32.98, p<0.0001; Dunnet’s multiple comparison test *p<.05, ****p<0.0001). Error bars = SEM.

Across all lines, most *C9orf72* mRNAs contained exon 1B, while exon 1A-containing transcripts represented only a small proportion of total transcripts (**Fig. 2c,d**). However, the sum of 1A and 1B transcripts was inferior to the total amount of *C9orf72* transcripts in lines containing the repeat expansion (C9-unedited, HET(WT)x) but not in mutation-corrected patient lines (REx, HET(Mut)x, 1Ax) (**Fig. 2c, Extended Data Fig. 12e**) or any of the WT lines (**Fig 2d, Extended Data Fig. 12f**). We hypothesize this gap corresponds to sense 1A-transcripts retaining the repeat expansion, onto which the primers would anneal too far apart for successful PCR. If so, the actual proportion of sense 1A-transcripts could reach 30% or more of total *C9orf72* transcripts in lines with the repeat expansion, up from <1% in corrected and WT cells (**Fig. 2c,d**). This does not include antisense repeat-containing transcripts, which are known to occur but are not captured by our assay, and would further increase the total amount of repeat-containing transcript. Interestingly, the amount 1A-transcripts decreased in all therapeutically edited C9 lines but not in the C9-HET(WT)x retaining the repeat expansion (**Fig. 2c, Extended Data Fig 12c**). Furthermore, there was no change in 1A-transcript levels between WT-unedited and lines in which the repeat-expansion had been removed (**Fig. 2d, Extended Data Fig 12d**). Together, these data suggest that 1A RNA transcription is upregulated in the diseased state.

To determine the proportion of transcripts coming from the WT and mutant alleles, we took advantage of a coding SNP (rs10757668) in the exon 2 splice acceptor of our patient line and phased it to the repeat expansion using single-molecule sequencing. Using probes that differed by a single nucleotide targeting this SNP, we determined what fraction of 1A- and 1B-transcripts derived from each allele (**Fig. 2e,f**). Once again and as expected, we could not detect 1A-transcripts containing the repeat expansion, presumably because of amplification failure. Nevertheless, most (>90%) exon 1A-containing transcripts came from the mutant rather than the WT allele in the unedited patient lines (**Fig. 2e**). The imbalance was corrected by repeat expansion excision. This finding suggests that at least some of the mutant 1A-transcripts can undergo normal splicing, and that the excess activity of 1A in the diseased state can be attributed for the most part to the mutant allele. Overall, these observations indicate an upregulation of exon 1A-transcripts off of the mutant allele, which implicates transcriptional upregulation of the mutation as a possible biological driver of disease.

In contrast to exon 1A transcripts, 1B-transcripts came predominantly (>68%) from the WT allele. Balanced bi-allelic expression of 1B-transcripts was restored after excision of the repeat expansion or simple excision of exon 1A (**Fig. 2f**). The latter finding indicates that the mere presence of the repeat expansion in the DNA does not cause the reduced 1B-transcript expression observed in the unedited line—enhanced activity of the 1A transcription start is a more likely explanation. As expected, excision of either allele resulted in elimination of expression from that allele (**Fig. 2e,f**).

### Allele-specific and bi-allelic excisions preserve production of normal levels of full-length C9orf72 protein

We quantified C9orf72 protein using the Simple Western system (WES). We validated antibody specificity using our knock-out line. Since exons 1A and 1B are non-coding, transcripts with either exon produce the same protein. None of the edits (REx, HET(Mut)x, HET(WT)x, 1Ax) reduced the C9orf72 protein levels in the patient line (**Fig. 2g,i**), and only exon 1A-excision reduced C9orf72 expression in the WT line (**Fig. 2h,j**). These results indicate that even major alterations of the *C9orf72* locus (such as removal of an entire allele) do not alter total protein levels in cells, which is advantageous for gene therapy.

We were interested in understanding the spatial localization of the C9orf72 protein in our edited lines, but unfortunately, after testing nine commercially available antibodies for immunocytochemistry, we found none that were specific to C9orf72 (*i.e*., they either had no signal or showed signal in our 2 KO lines) (**Supplementary Table 4, Extended Data Fig. 13**).

### Bi-allelic and allele-specific excisions abolish expression of repeat-encoded polypeptides

*C9orf72* is transcribed off of both the sense and anti-sense strands, in both normal and diseased cells (**Fig. 3a**)^20, 28, 74, 75^. Our data suggests that sense transcription of the repeat region starts from exon 1A, since excision of exon 1A closed the gap in “undetectable” sense transcript (**Fig. 2c, Extended Data Fig. 12c,e**). However, it is unknown where anti-sense transcription initiates. The repeat expansion is translated through non-canonical RAN translation from transcripts derived from both and sense and anti-sense strands to form five DPRs which are thought to be toxic^19, 20, 23^ (**Fig. 3a, Extended Data Fig. 14a**). We therefore evaluated the ability of therapeutic edits to prevent the production of DPRs. To this end, we measured two DPRs, one (poly-GA) encoded exclusively on the sense strand, the other (poly-GP) on both strands (**Fig. 3a**) in our C9-isogenic series. We evaluated ten antibodies targeting each of these DPRs using MSD’s sandwich ELISA and found two antibody combinations that could reliably detect the presence of poly-GA and poly-GP DPRs above the background level defined by our KO line (**Extended Data Fig. 14b**). Poly-GA expression was eliminated by each of the therapeutic edits (C9-REx, HET(Mut)x, 1Ax) but unchanged by excision of the WT allele (C9-HET(WT)x) (**Fig. 3b**). Poly-GP was eliminated by removal of the repeat expansion (REx, HET(Mut)x) but not by excision of exon 1A (**Fig 3c,** 1Ax), consistent with the proposal that it also derives from anti-sense transcription of the repeat expansion. Excision of the WT allele more than doubled the amount of poly-GP expression (**Fig. 3c,** HET(WT)x), indicating an interaction between alleles that is worthy of further exploration.

**Figure 3.**
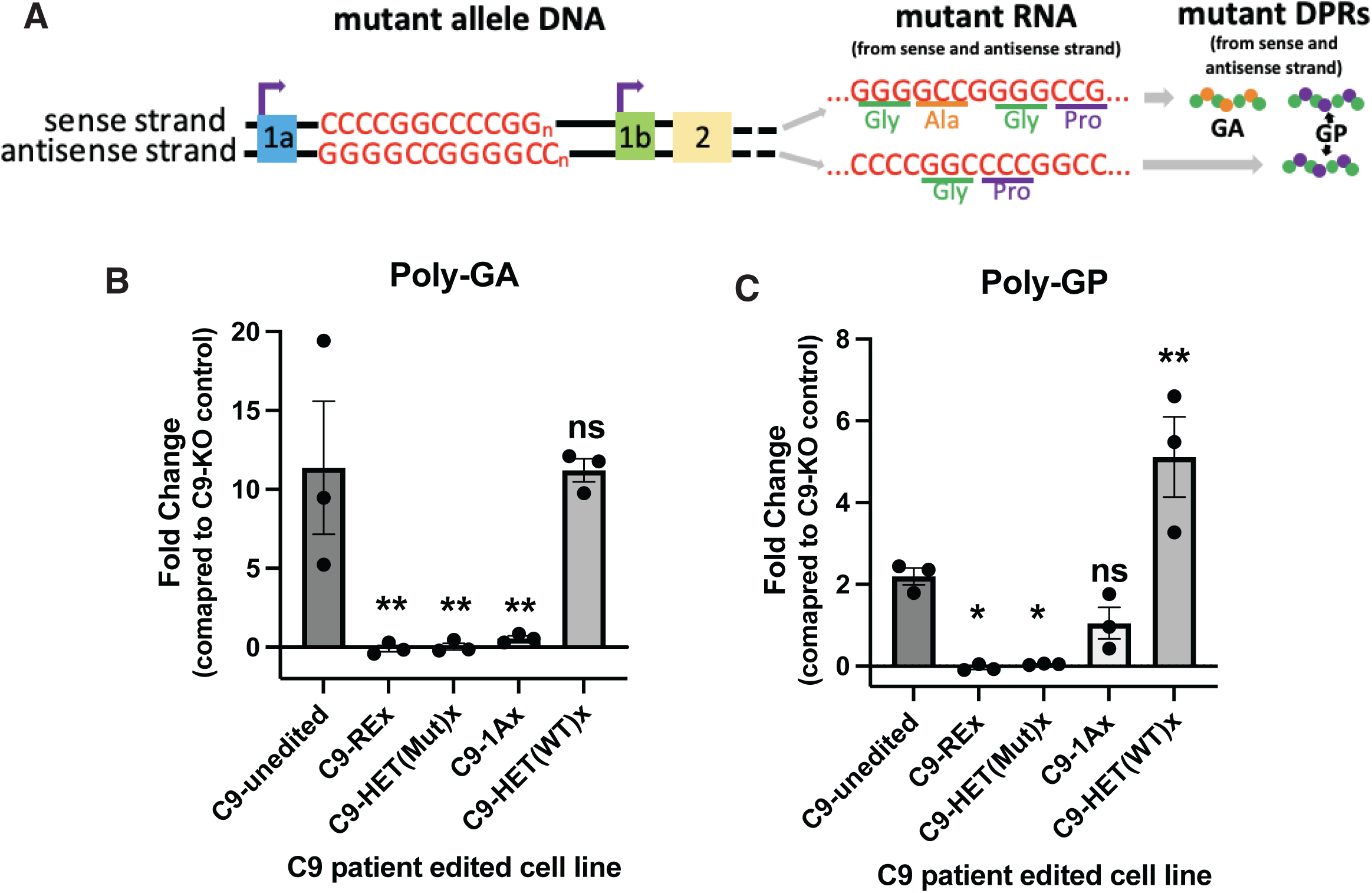
Sense dipeptide repeat expression (DPR) is corrected by three therapeutic gene editing approaches whereas antisense DPR expression is only corrected by removing the repeat expansion. (A) The repeat expansion is transcribed from the sense strand (starting at exon 1A) and the antisense strand and gives rise to polyGA and polyGP peptides through non-canonical repeat-associated non-AUG (RAN) translation. Only sense transcription can give rise to a Poly-GA peptide, whereas Poly-GP can arise from the sense or antisense strands. (B,C) Quantification of Poly-GA (B) and Poly-GP (C) in C9-unedited and edited cell lines above baseline noise established by the C9 KO line, as measured via MSD sandwich ELISA. Only excision of the repeat expansion or the mutant allele abolishes production of both Poly-GA (1-way ANOVA F(4,10)=10.12, p<0.001) and Poly-GP peptides (1-way ANOVA F(4,10)=19.66, p<0.0001). Excision of exon 1A only abolishes expression of Poly-GA, consistent with silencing of sense but not anti-sense transcription. Excision of the WT allele (HET(WTx)) more than doubled expression of Poly-GP. *p<0.5, **p<0.01 by Dunnet’s multiple comparisons test. Error bars = SEM.

### Bi-allelic and allele-specific excisions revert TDP-43 pathology in 7-week-old patient-derived neurons

The pathological hallmark of C9FTD/ALS is loss of nuclear TDP-43 and TDP-43 aggregation in the cytoplasm of affected neurons^76^. These events are thought to be independent^76^ and have been difficult to model in cellular or animal systems. We detected a clear loss of nuclear TDP-43 in 57% of TDP-43-positive neurons derived from our unedited patient cell line (**Fig. 4a, pink arrow**). By contrast, this rate was on average less than 30% in each of the therapeutically edited lines (REx, HET(Mut)x, 1Ax). Loss of nuclear TDP-43 was only apparent after aging neurons in culture for 7-weeks post-differentiation, suggesting an interaction between aging and genotype.

**Figure 4.**
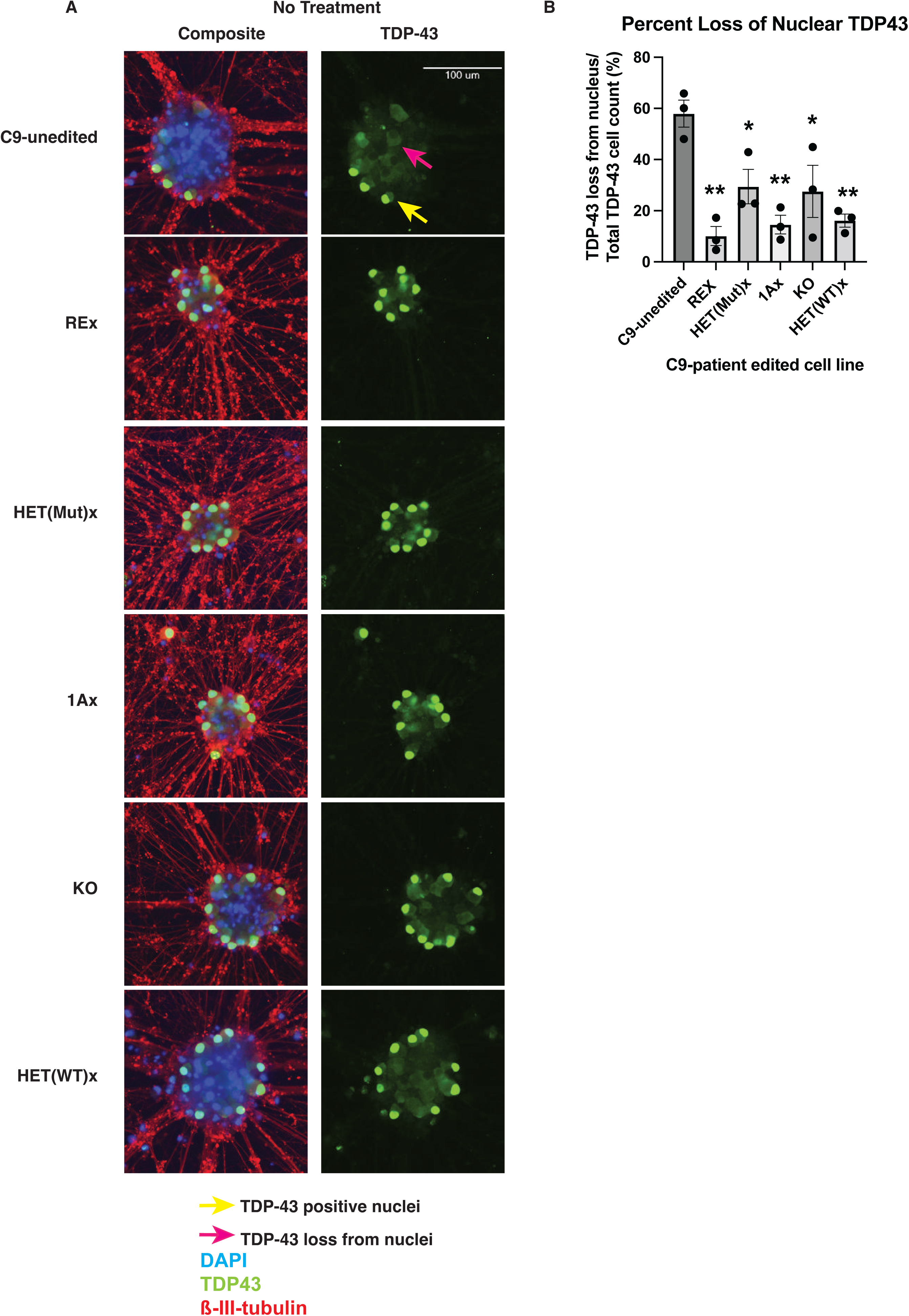
Three editing approaches correct loss of nuclear TDP-43 in 7-week-old neurons derived from C9 line. (A) Immunofluorescent images of neurons derived from unedited or edited C9 iPSCs. The neurons were grown for 7 weeks and stained for TDP-43 (green), DAPI (blue) and beta-III-tubulin (red). Yellow arrow points to a nucleus harboring TDP-43 and pink arrow to a TDP-43-positive cell whose nucleus is devoid of TDP-43. (B) Percentage of TDP-43- positive cells that lack nuclear TDP-43 (1-way ANOVA F(5,12)=8.61; *p<0.05, **p<0.01 by Dunnett’s multiple comparison’s post-hoc test). Each experiment contained 3 biologic replicates (separate wells). Error bars = SEM.

## Discussion

Up to now, the *C9orf72* locus has been challenging to edit in a manner compatible with clinical translation. We investigated three strategies we thought had therapeutic potential because they did not involve template-based gene correction, which is inefficient in post-mitotic neurons, and because they minimized off-target editing. Each strategy capitalized on Cas9’s ability to cut DNA, which aligns with technologies that are closest to clinical prime-time^63, 64^.

We found that two of our three approaches (excision of the repeat region and excision of the mutant allele) corrected RNA abnormalities, preserved protein levels, and reverted dipeptide repeat and TDP43 pathology in iPSC-derived neurons from a patient line harboring ∼200 repeats. By contrast, our attempt at silencing rather than removing the repeat expansion proved suboptimal, as it only blocked sense transcription and did not eliminate the production of toxic peptides from anti-sense transcripts. From these data, we can advance repeat expansion excision and allele-specific excision to further pre-clinical testing. To determine which of these two approaches is more efficient and precise, we will need to edit in post-mitotic neurons directly as well as *in vivo* models. Our current findings suggest that excision of the repeat expansion is not very efficient in diseased iPSCs, in contrast to the much larger excision of the mutant allele. These findings warrant further investigation across patient lines with various repeat lengths, and, ultimately, in differentiated patient-derived neurons.

While a significant step forward in developing gene editing for neurodegenerative diseases, this work would be greatly enhanced by technological advancements in CRISPR delivery technology for *in vitro* and *in vivo* gene editing. Viral vectors are currently used for *in vitro* delivery to neurons. However they risk increasing off-target rates^77, 78^ given their continual expression of Cas9/gRNA. They also risk unwanted incorporation of viral DNA^79, 80^. We are eager to develop and adopt delivery technologies for additional pre-clinical testing to advance toward clinical trials. This would also allow us to rigorously test on- and off-target rates in the disease-relevant cell type that could help us differentiate between our alternate editing strategies.

Our robust editing and outcome measurement tools lay the groundwork to investigate gene-editing approaches for monogenic disease in human iPSCs and derived cell-types relevant to disease, and are applicable to any monogenic disease, particularly other repeat expansion disorders. In particular, we demonstrated the usefulness and reliability of single-molecule sequencing to characterize large repeat expansions and verify their excision, and we recommend that this approach become the gold standard for future studies of repeat expansion diseases. While our work generated nine engineered lines across a patient and control background to establish these methods, it will be important to reproduce our findings across other patient lines, especially those with different repeat expansion sizes. While loci with expanded repeats each have their idiosyncrasy, our work offers lessons that should apply beyond *C9orf72*. One is that several gene editing approaches should be compared side by side in patient lines, as we were surprised to find that a 22 kb allele-specific excision was more efficient than a 227 bp excision. Another lesson is that a detailed understanding of the organization of the target gene locus and expression is fundamental, to ensure that unexpected side effects, such as amplifying DPR expression by excising the WT allele, are adequately detected. Perhaps the greatest lesson is that the best approach remains to remove the repeat expansion itself, as expanded repeats generate expression patterns that are difficult to anticipate and revert, and complete neutralization of the mutant allele may not be well tolerated at loci other than *C9orf72*.

---

**Extended Data Figure 1.**
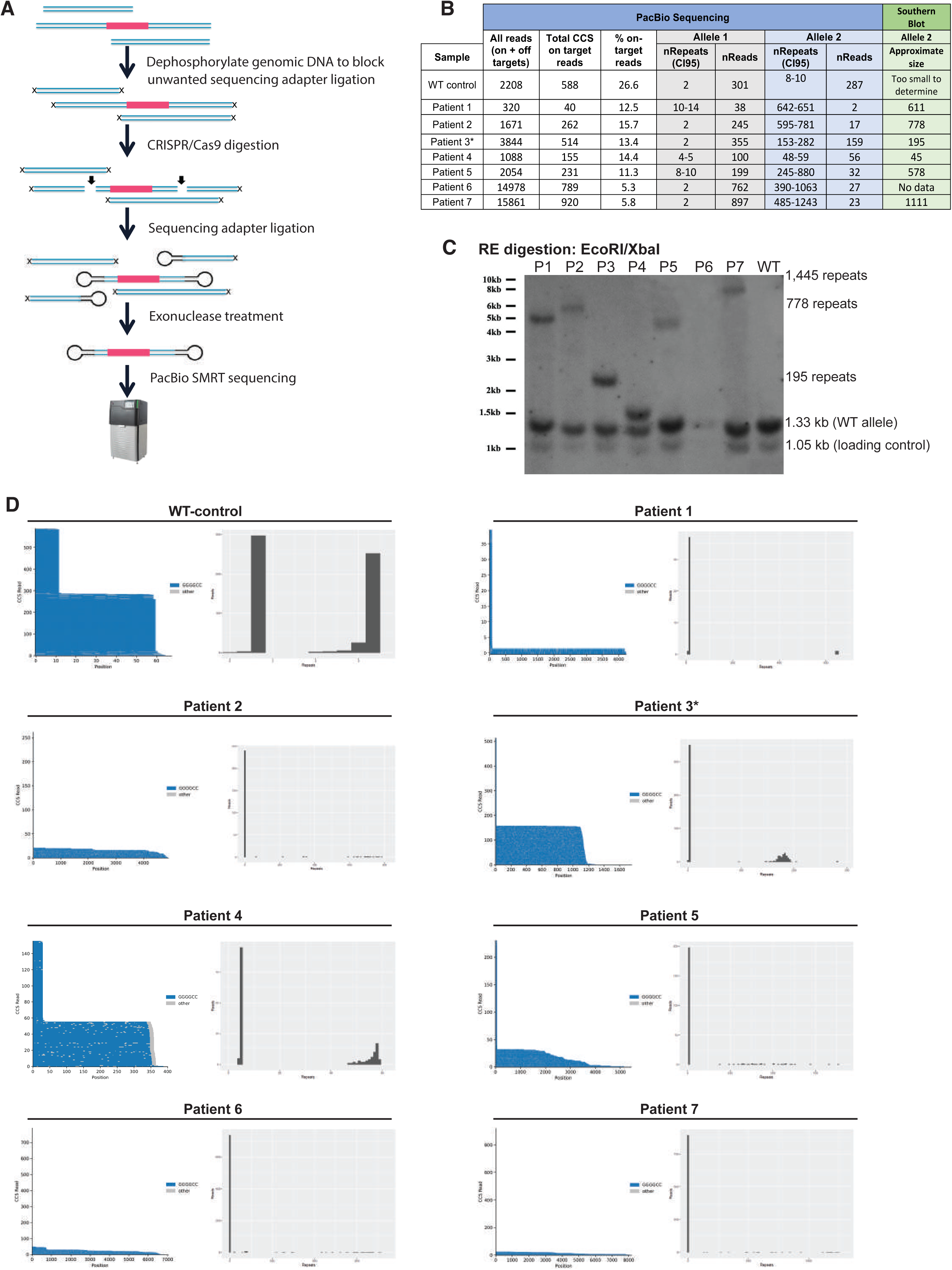
Pacific Biosciences (PacBio) single molecule sequencing to determine the repeat size in 8 iPSC lines. (A) Schematic of the pipeline used to generate the library for single molecule sequencing. We excised the repeat region (red) from high molecular weight DNA using CRISPR and guide RNAs flanking the repeat regions (arrows). We then seal the CRISPR-generated double-strand breaks by ligating in sequencing adapters. Subsequent exonuclease treatment results in an enrichment for the excised repeat region, which is sealed at both ends. The enriched repeat regions are then subjected to PacBio SMRT sequencing. Because the sequenced molecules are circular, the sequencing reaction can read through them more than once, which increases the accuracy of the sequencing data. Barcoding allows us to multiplex samples to reduce sequencing costs. (B) We sequenced 3-5 µg of DNA from 1 WT-control iPSC line and 7 iPSC lines from patients harboring expansions of the *C9orf72* repeat. Allele-specific SNPs allowed us to distinguish the repeat regions from both *C9orf72* alleles (Allele 1 and Allele 2) in each cell line. On-target reads are reads that sequenced the entire excised region (including the repeat region and flanking DNA) 3 times (= 3 pass criteria). Within these reads, we counted the number of GGCCCC repeats starting right after an anchor (CGCCC) 5’ to the repeat region. Repeat lengths and associated read counts are reported for each allele of each cell line and compared to repeat length estimated by Southern blot. Repeat lengths estimated by Southern blot were comparable to mean repeat lengths determined by single-molecule PacBio sequencing. (C) Southern blot of nuclear DNA from WT control and patient iPSCs listed in (B). After EcoR/XbaI digestion, a loading control fragment (1.05kb), WT fragment (1.33kb) and fragments with repeat expansions of various lengths were detected. Southern blot required 20 µg of input DNA (vs. 3-5 µg input for PacBio sequencing) and a sample with 14 µg (P6) failed detection, demonstrating the insensitivity of Southern blot. (D) Sequencing traces (left graphs) and histograms showing the number of CCS reads per repeat count (right graphs). In the sequencing traces, each horizontal line represents one sequenced molecule of DNA. Blue color depicts on target sequencing, grey color depicts sequencing error. Each molecule is anchored to an adjacent, non-repeat region (CGCCC) which is not included in the total repeat count. Y-axis = circular consensus sequencing (CCS) count. In the WT line, repeat length has a bimodal distribution with roughly equal numbers of reads containing 2 or 10 repeats, indicating that one allele has 2 repeats and the other between 8 and 10. In the patient lines, a bimodal distribution is not always as apparent: all lines show a peak with a low repeat number (2-10), corresponding to the unexpanded allele, but the expanded allele can yield a wide range of repeat counts. The iPSC line from patient 3 (P3*), with a bimodal distribution centered at 2 and 175, was selected for our excision studies.

**Supplementary Table 1.**
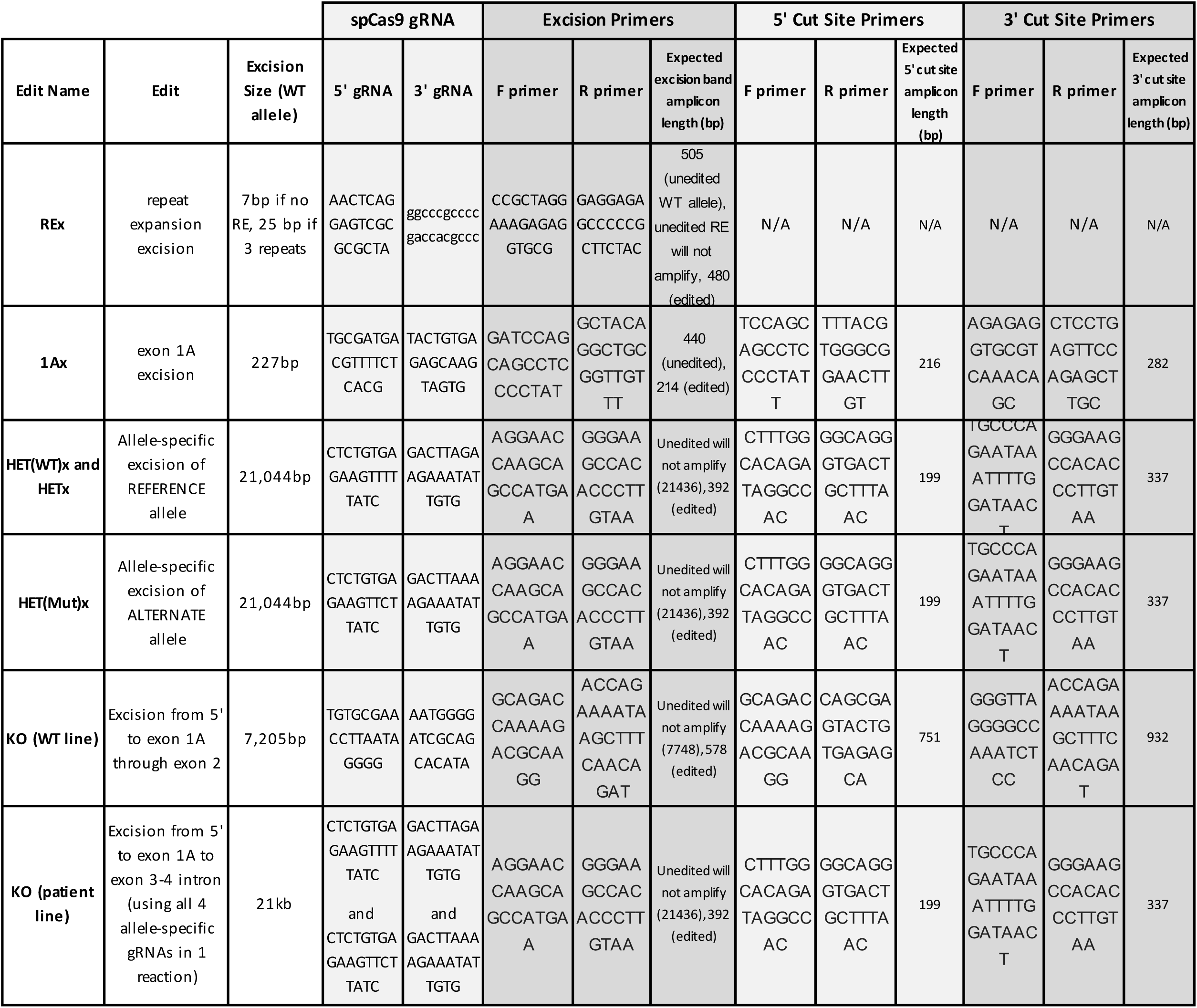
**Guide RNAs and primers** used to generate and verify, respectively, each edited cell line. Excision size is provided for type of excision in the WT line. Expected amplicon size for each set of PCR primers is also provided.

**Supplementary Table 2.**
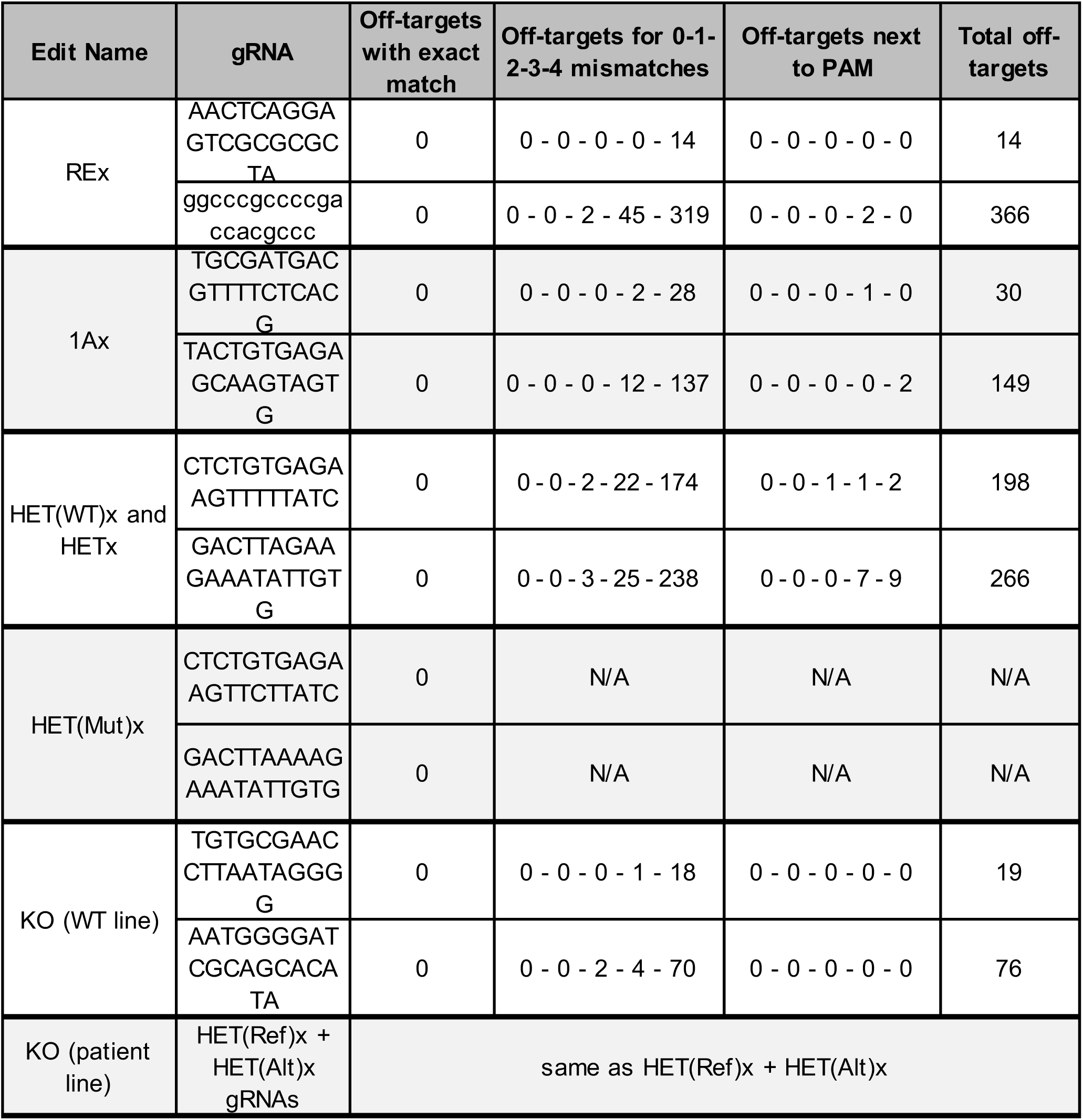
**Predicted off-targets for each gRNA.** We used CRISPOR (Homo sapiens – USCS Dec. 2013 (GRCh38/hg38)) to predict off-targets for each gRNA combined with spCas9. Off-targets are displayed as a function of the number of mismatches (0-1-2-3-4) in the entire 20 nucleotide gRNA sequence. For example REX 3’ gRNA has 0-0-2-45-319 predicted off-targets meaning 0 off-targets with 0 or 1 mismatches, 2 off-targets with 2 mismatches, 45 off-targets with 3 mismatches and 319 off-targets with 4 mismatches. Off-targets next to the PAM indicates predicted off-targets that have no mismatches in the first 12 bp adjacent to the PAM. These are more likely to be true off-targets. CRISPOR only considers off-targets adjacent to an NGG, NAG, NGA PAM.

**Extended Data Figure 2.**
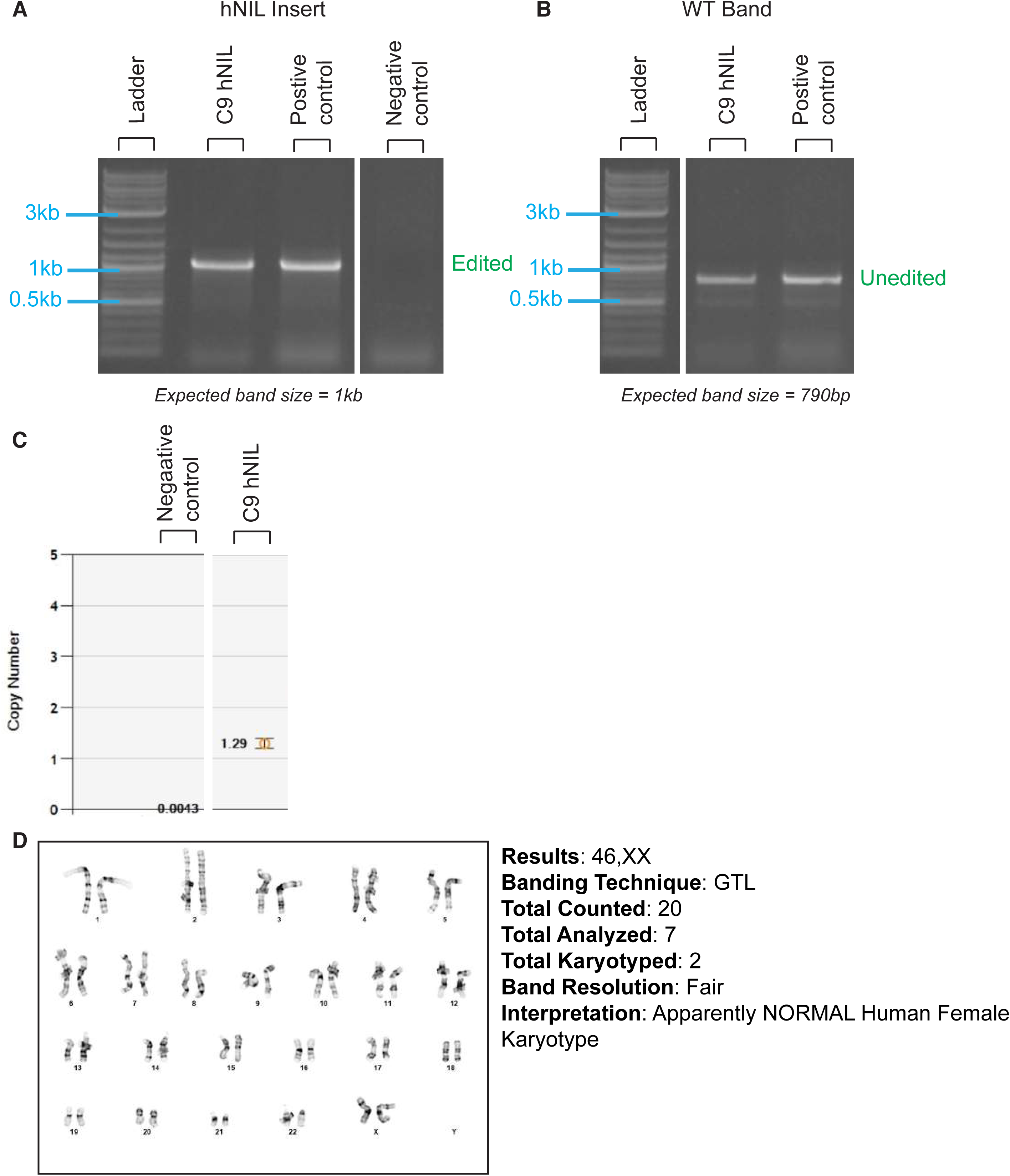
Construction of C9-control (unedited) hNIL cell line. (A) Band corresponding to the insertion of the hNIL construct into an unedited patient cell line (also called C9-unedited), as determined by PCR. (B) Preservation of a WT band indicates the line is heterozygous. (C) ddPCR copy number assay shows 1 insertion of the hNIL construct. (D) The cell line had a normal karyotype.

**Extended Data Figure 3.**
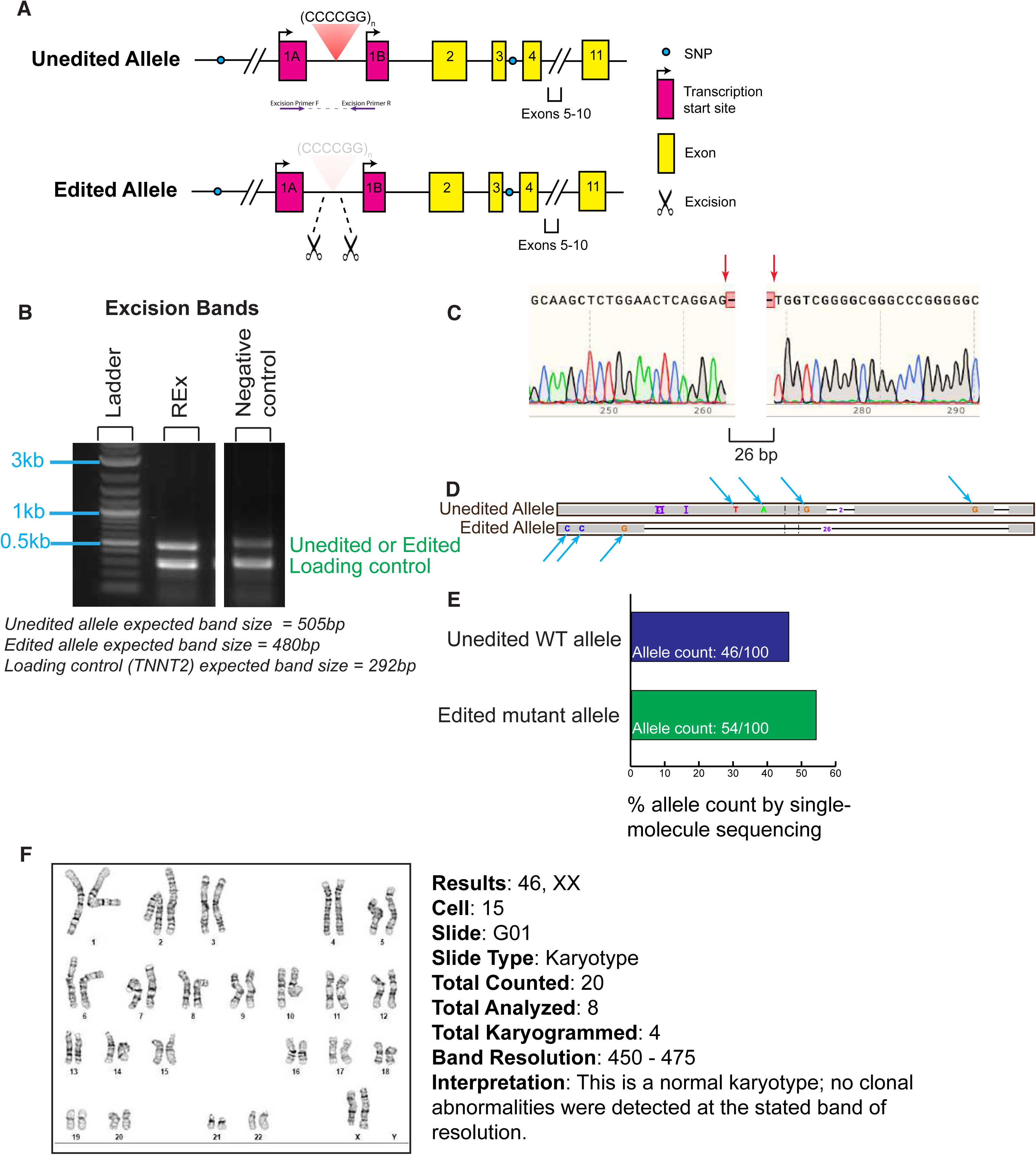
Construction of the C9-REx cell line. (A) Position of the gRNAs (indicated by scissors) and excision primers (purple arrows) used to create and verify, respectively, excision of the repeat expansion (REx) in the *C9orf72* gene in a patient cell line. (B) Line had a band at ∼500bp using excision primers and clean Sanger sequencing (C; cut sites indicated by pink arrows). However, we could not tell from these data whether the line had a homozygous excision of the repeat region or a heterozygous excision of the WT allele only, since the expanded repeat region fails amplification. We therefore used single-molecule sequencing (D) to determine that the clone was pure, and carried a heterozygous excision of the expanded repeat leaving a 26 bp deletion on the mutant allele (using SNPs to differentiate alleles, indicated by blue arrows). (E) Allele count of PacBio sequencing data shows both alleles were equally covered by sequencing. (F) The cell line had a normal karyotype.

**Extended Data Figure 4.**
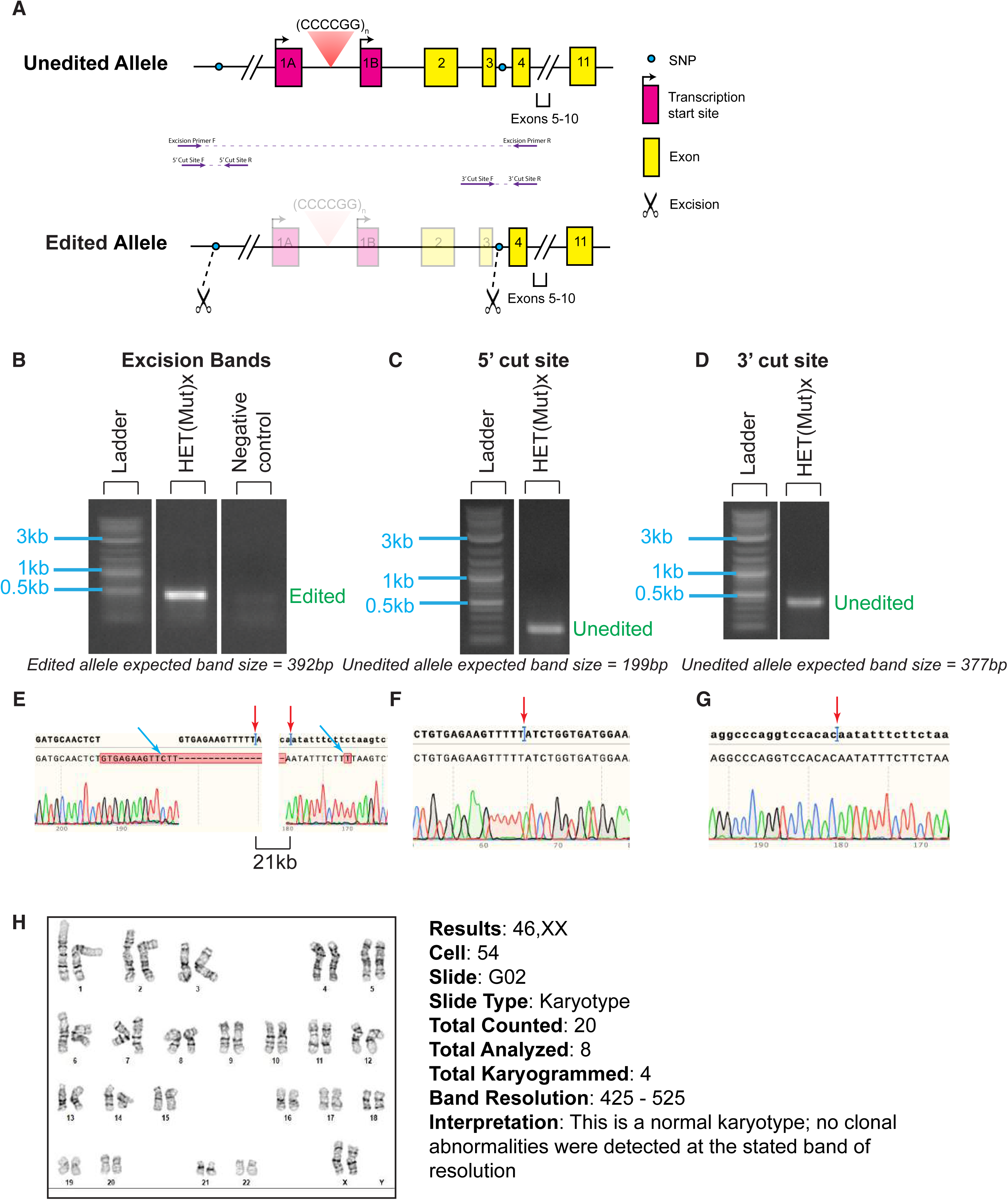
Construction of the C9-HET(Mut)x cell line. (A) Position of the gRNAs (indicated by scissors) and excision and cut site primers (purple arrows) used to create and verify, respectively, a ∼22kb excision of the mutant *C9orf72* allele in a patient cell line. SNPs phased to the repeat expansion (blue dots) were used to target the mutant allele. Presence of an excision band (B) and preservation of bands at both the 5’ (C) and 3’ (D) cut sites indicates the line is a heterozygous excision. Corresponding clean Sanger sequencing (D-G) shows the clone is pure. (pink arrow – cut site; blue arrow – misaligned Sanger sequencing). (H) The cell line had a normal karyotype.

**Extended Data Figure 5.**
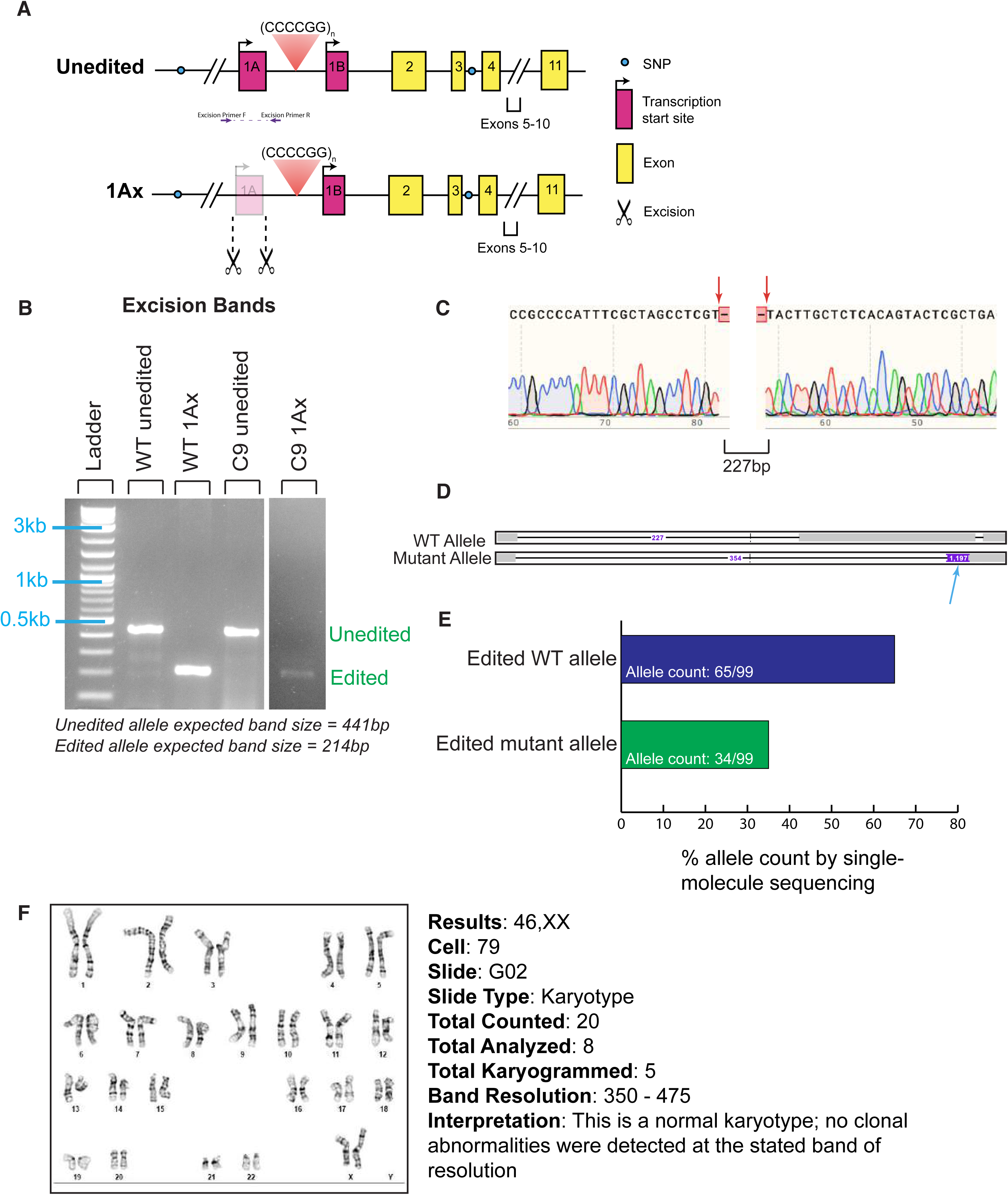
Construction of the C9-1Ax cell line. (A) Position of the gRNAs (indicated by scissors) and excision primers (purple arrows) used to create and verify, respectively, excision of exon 1A of the *C9orf72* gene in a patient cell line. (B) Presence of an excision band and absence of a WT band C9-1Ax indicates the line is homozygous. WT-unedited, C9-unedited and WT-1Ax serve as negative and positive controls. (C) Sanger sequencing shows the excision cut sites (pink arrows). (D) Single-molecule sequencing revealed 227 bp excision on the WT allele and a 354 bp excision on the mutant allele (blue arrow shows the repeat expansion). (E) Total alleles sequenced by single molecule sequencing showed a modest preference for the WT allele, as expected. (F) The cell line had a normal karyotype.

**Extended Data Figure 6.**
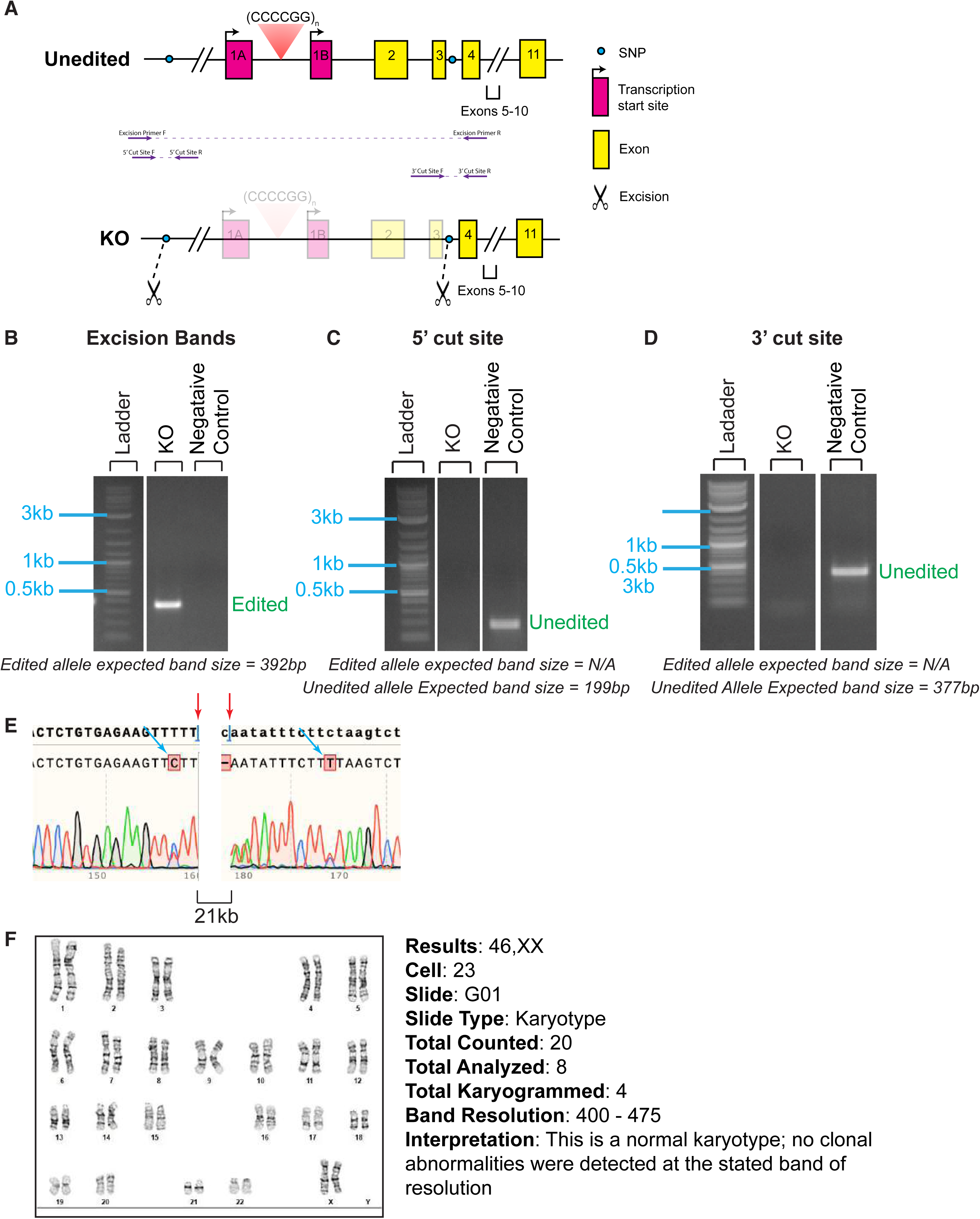
Construction of the C9-KO cell line. (A) Position of the gRNAs (indicated by scissors) and excision and cut site primers (purple arrows) used to create and verify, respectively, a ∼21 kb excision of both the mutant and WT *C9orf72* alleles in a patient cell line. Line was made by using 4 allele-specific gRNAs targeting SNPs (blue circle) in one reaction. (B) Presence of an excision band and absence of WT cut site bands (C,D) indicate homozygosity. (E) Sanger sequencing shows the excision cut sites (pink arrows). (F) The cell line had a normal karyotype.

**Extended Data Figure 7.**
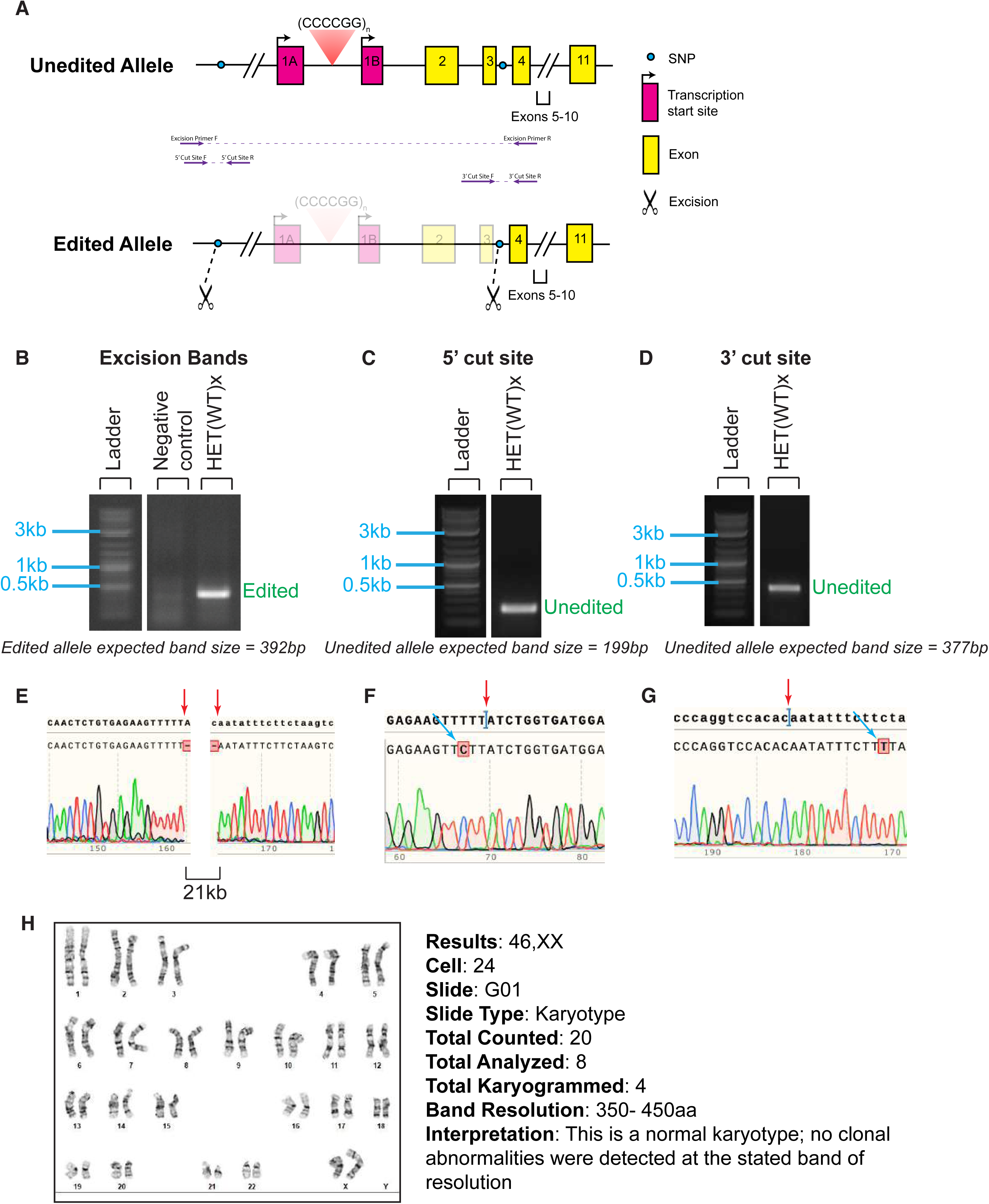
Construction of the C9-HET(WT)x cell line. (A) Position of the gRNAs (indicated by scissors) and excision and cut site primers (purple arrows) used to create and verify, respectively, excision of the WT *C9orf72* allele in a patient cell line. The SNPs phased to the WT allele (blue dots) were targeted to create the 21 kb excision. Presence of an excision band (B) and preservation of bands at both the 5’ (C) and 3’ (D) cut sites indicate the line has a heterozygous excision. Corresponding clean Sanger sequencing (D-G) shows the clone is pure. (pink arrow – cut site; blue arrow – SNPs). (H) The cell line had a normal karyotype.

**Extended Data Figure 8.**
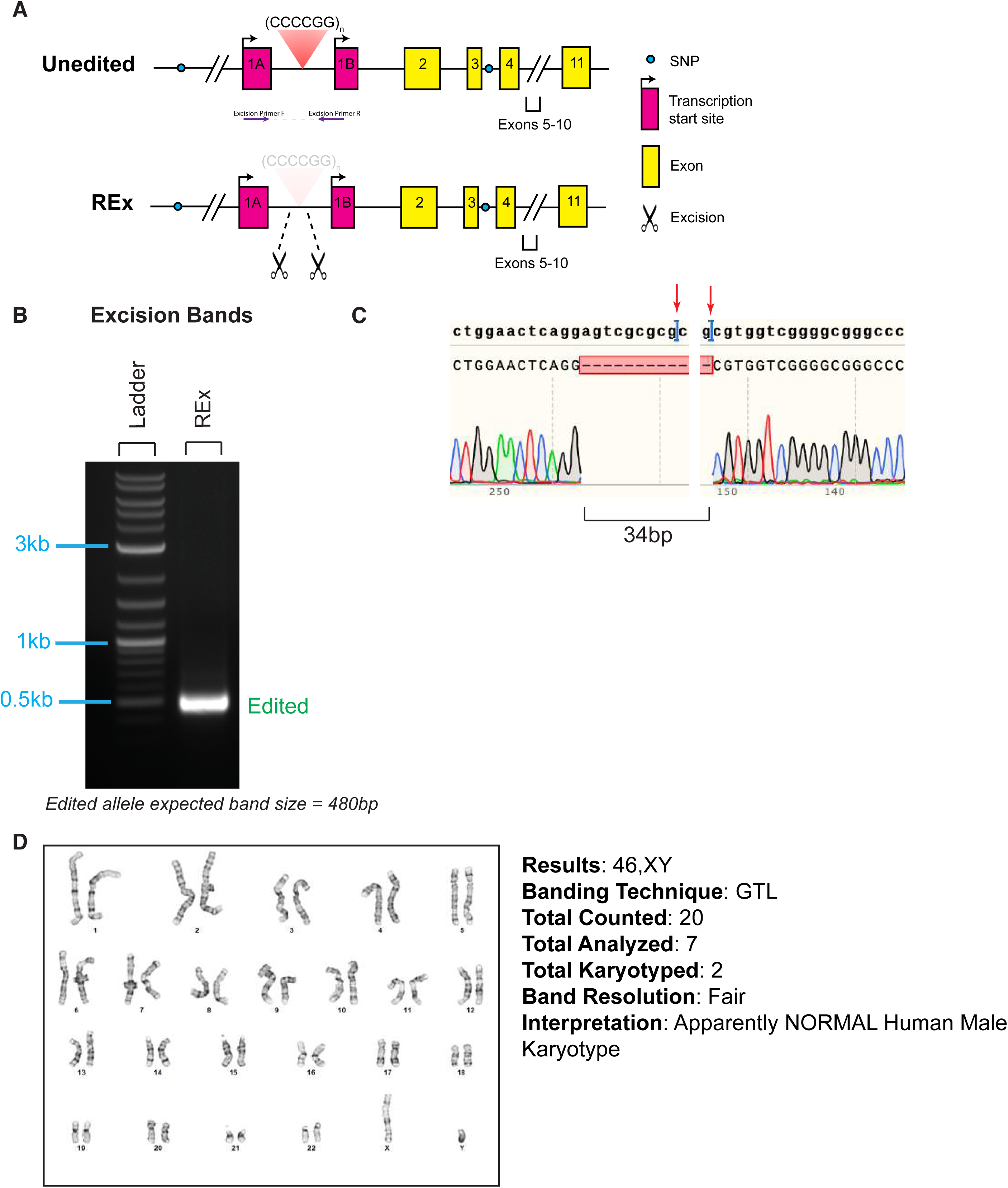
Construction of the WT-REx cell line. (A) Position of the gRNAs (indicated by scissors) and excision primers (purple arrows) used to create and verify, respectively, excision of the repeat region of the *C9orf72* gene in a non-diseased (control) cell line. (B) Presence of an excision band and Sanger sequencing (C) show the 34 bp excision. We could not perform cut site sequencing because we could not design unique primers inside the GC-rich repeat region. Therefore we relied on clean Sanger sequencing (C) to indicate the line was pure around the cut sites (pink arrows). (D) The cell line had a normal karyotype.

**Extended Data Figure 9.**
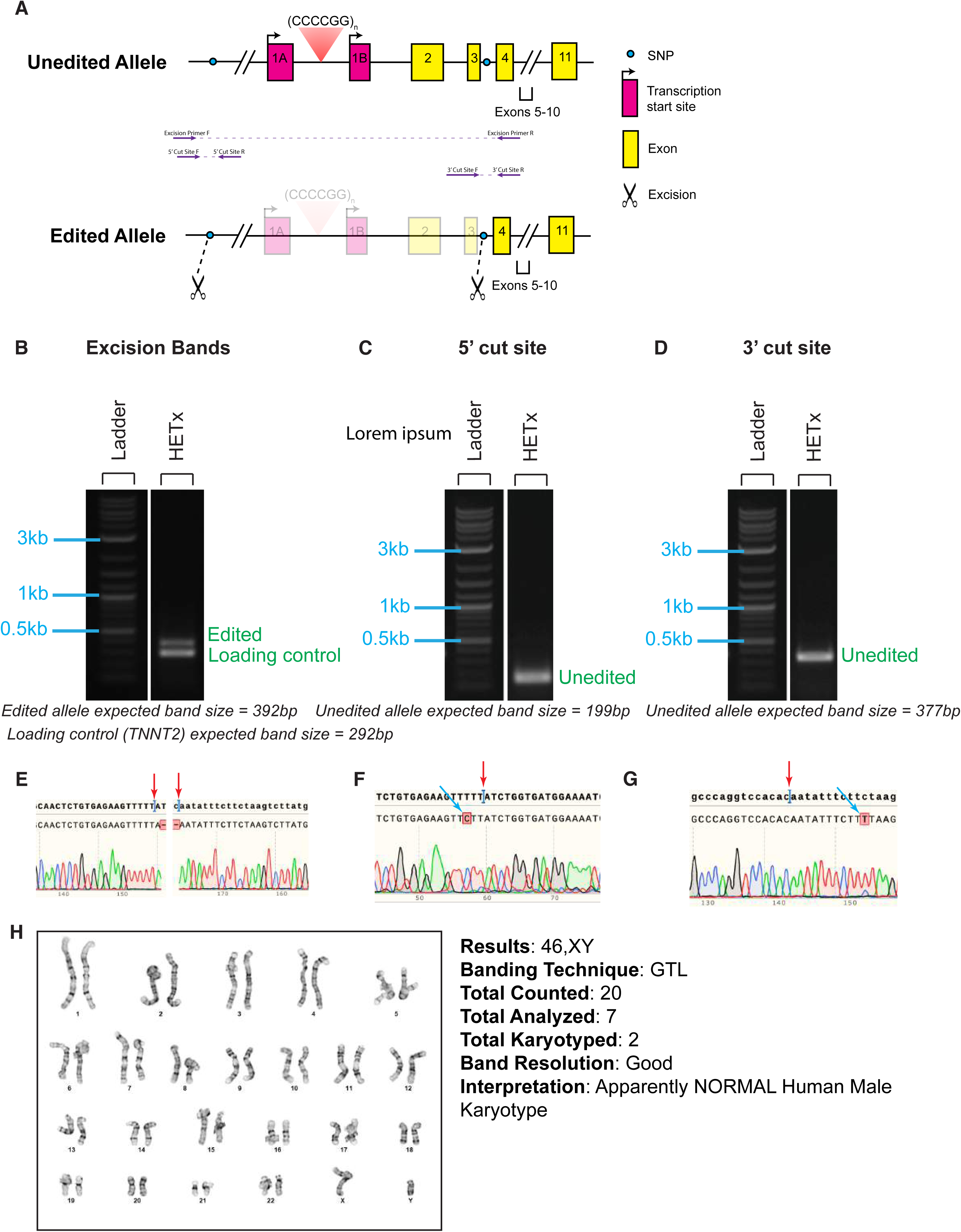
Construction of the WT-HETx cell line. (A) Position of the gRNAs (indicated by scissors) and excision primers (purple arrows) used to create and verify, respectively, excision of the repeat region of the *C9orf72* gene in a non-diseased (control) cell line. SNPs phased to the repeat region (blue dots) were used to target a single allele. Presence of an excision band (B) and preservation of bands at both the 5’ (C) and 3’ (D) cut sites indicate the line has a heterozygous excision. Corresponding clean Sanger sequencing (D-G) shows the clone is pure. (Pink arrow – cut site; blue arrow – SNP). (H) The cell line had a normal karyotype.

**Extended Data Figure 10.**
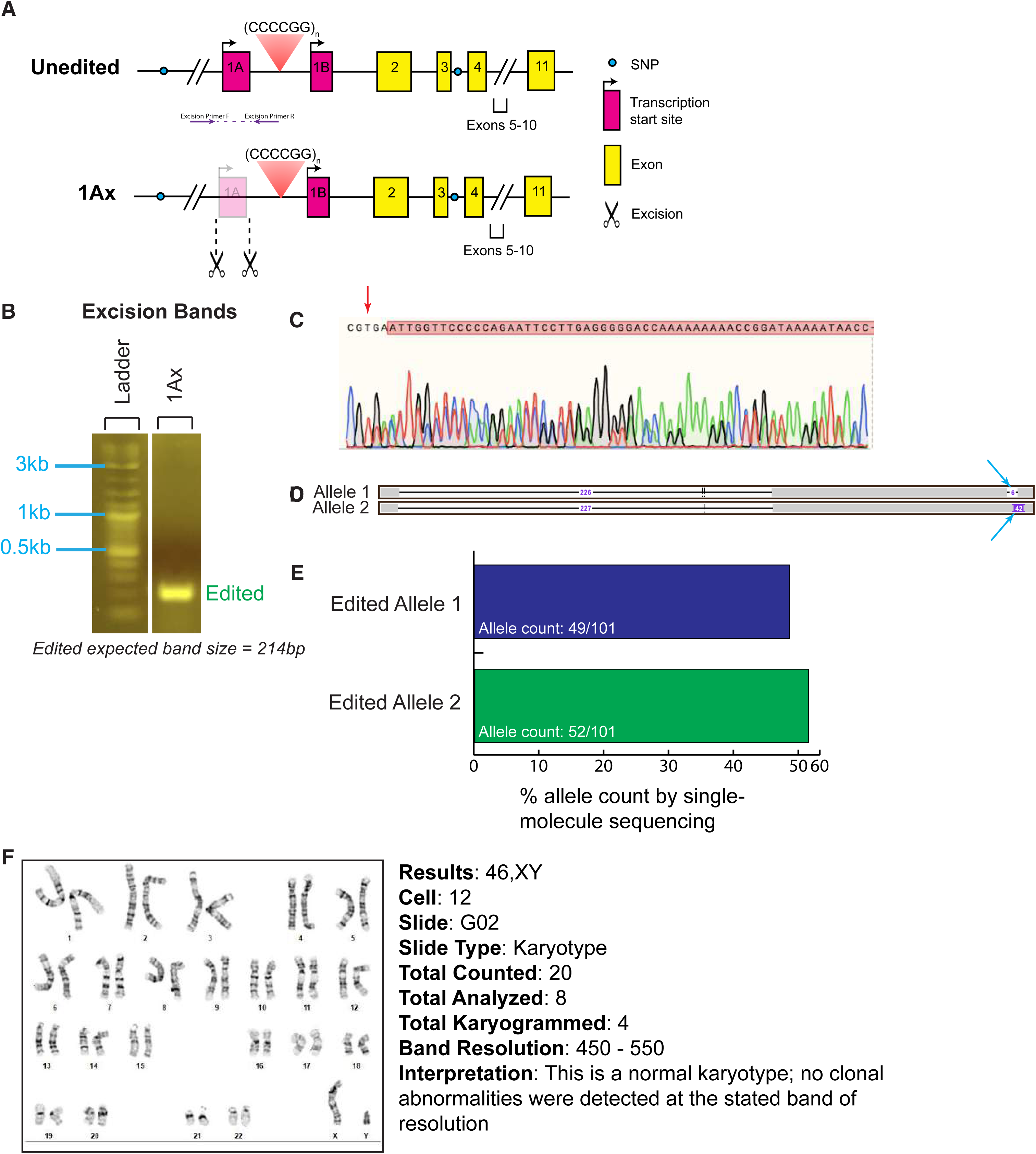
Construction of the WT-1Ax cell line. (A) Position of the gRNAs (indicated by scissors) and excision primers (purple arrows) used to create and verify, respectively, excision of exon 1Ax of the *C9orf72* gene in a non-diseased (control) cell line. (B) Excision band was present by PCR. (C) Because of messy Sanger sequencing after subcloning which appeared to show a 1 bp overlap in traces after the cut site (pink arrow), we could not resolve whether the clone remained impure or whether there were different editing outcomes on the two alleles. We therefore turned to single-molecule sequencing (D), which indicated a pure clone with a 226 bp excision on one allele and a 227 bp excision on the other. (E) The percentage of each allele detected by single-molecule sequencing was roughly equal. (F) The cell line had a normal karyotype.

**Extended Data Figure 11.**
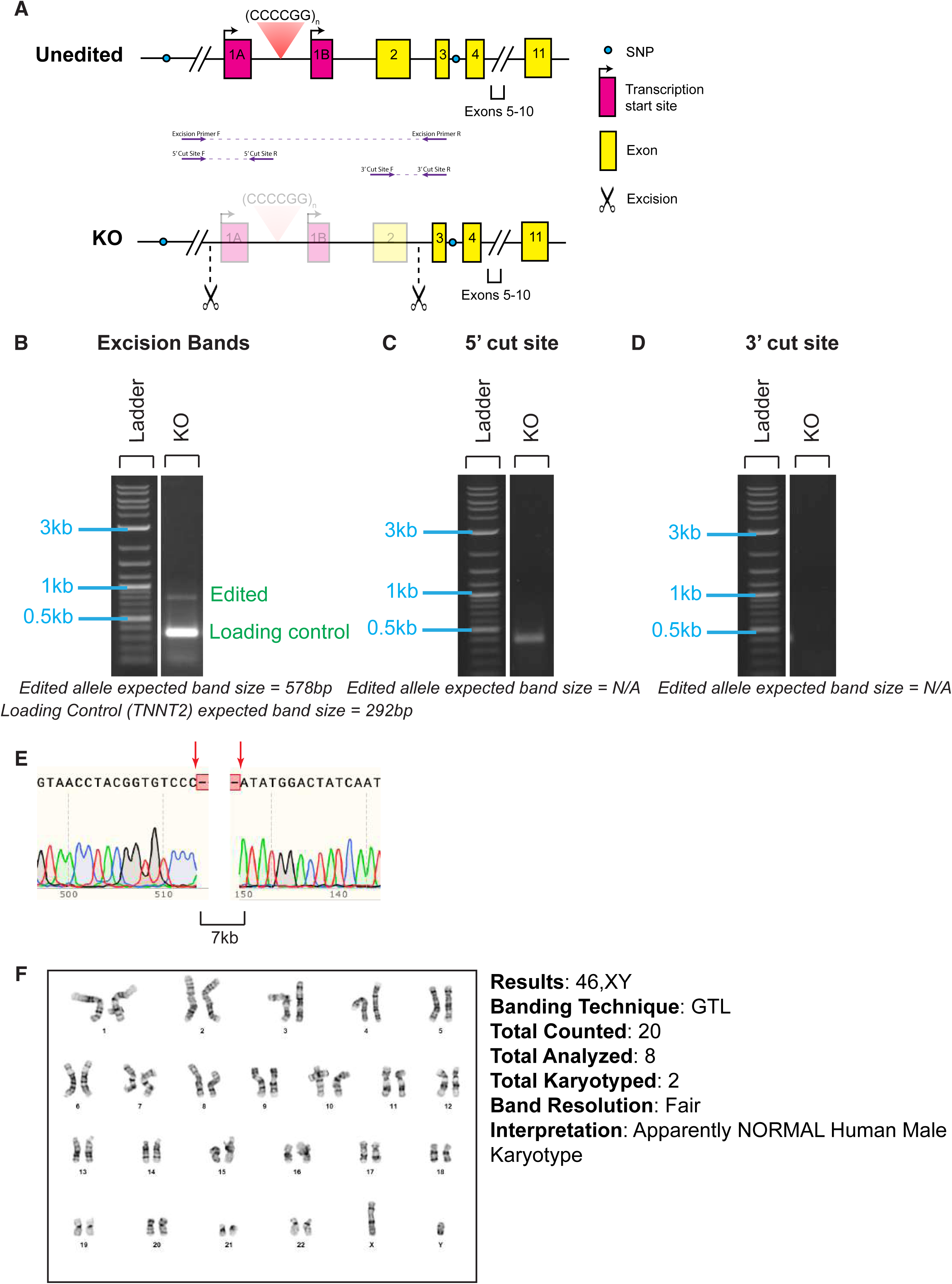
Construction of the WT-KO cell line. (A) Position of the gRNAs (indicated by scissors) and excision primers (purple arrows) used to create and verify, respectively, a 7 kb biallelic excision of the *C9orf72* gene in a non-diseased (control) cell line. (B) Excision band was 153 bp larger than the expected 578 bp, corresponding to a smaller than predicted excision. (C,D) Absence of WT cut site bands at 751 5’ and 932 3’ indicate homozygosity. We sequenced the unexpected 400 bp band at the 5’ cut site (C) which had no homology to the *C9orf72* locus, indicating a primer off-target. (E) Sanger sequencing shows the excision cut sites (pink arrows). (F) The cell line had a normal karyotype.

**Extended Data Figure 12.**
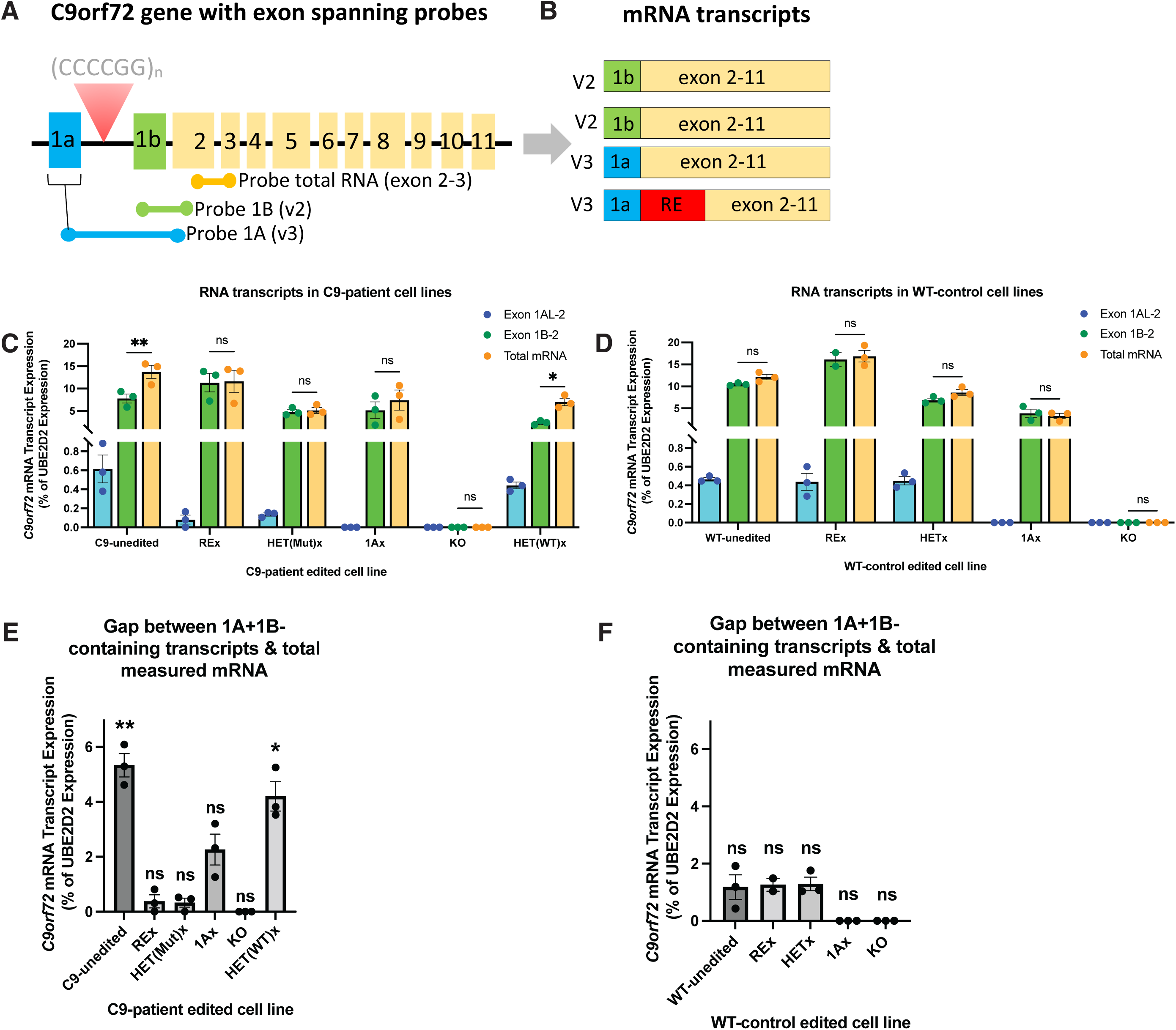
Quantification of C9orf72 RNA in edited C9-patient and WT-control lines. (A,B) We used exon-spanning PCR primers to quantify RNA variant incorporating exon 1A, exon 1B or total RNA (exon 2-3 spanning). (C,D) ddPCR quantification of exon 1A (V3) RNA (blue), exon 1B (V2) RNA (green) and total RNA (orange) in isogenic lines from a C9-patient (C) or WT-control (D). These data are depicted in Fig 2 C,D and repeated here to illustrate total contribution from each transcript variant. We detected a significant gap between exon-1B- containing transcript and total transcript that was only present in lines harboring a repeat expansion (C9-unedited, HET(WT)x) (mixed models F(10,36) = 5.6, p<0.0001; Tukey’s post-hoc test *p<0.05, **p<0.005). This gap was closed in C9-corrected lines (C9-REx, HET(Mut)x, 1Ax) and all WT lines (WT: mixed models F(8.29)=38.9, p<0.0001). (E,F) Quantification of the gap between detectable RNA variants (1A + 1B-containing transcripts) vs total measured RNA (exon 2-3 containing transcripts) in C9-patient (E) and WT (F) lines. Only C9-lines expressing the repeat expansion (C9-unedited, HET(WT)x) significantly differed from 0 (0 = no gap between measured variant and total transcript) (one-sample t-test corrected for multiple comparisons, *=p<0.01). We hypothesize that mutant sense RNA comprises the gap between total measured RNA and measured exon 1A+1B-transcripts since the presence of a repeat expansion would disrupt exon 1A-exon 2 primer amplification as well as probe binding.

**Supplementary Table 3.** Exon-spanning ddPCR probes used in Figure 2 and Extended Data Fig. 12.

| ddPCR Design | Probe target | Catalog Name | Vendor | Catalog Number | Fluorophore |
| --- | --- | --- | --- | --- | --- |
| 1A-long transcript (variant 3) | Exon 1A-exon 2 junction (NM_001256054.2) | Hs00948764_m1 for C9orf72 | Thermo | 4351372 | FAM |
| 1B Long | Exon 1B-exon 2 junction (NM_018325.4) | ID: AII1NCR, Name:C9_NEWV2 | Thermo | 4331348 | FAM |
| Exon 2-3 total | Exon 2-3 junction (total mRNA) | Hs00376619_m1 | Thermo | 4448892 | VIC |
| 1A Allele Specific | Exon 1A-exon 2 junction (NM_001256054.2) targeting SNP rs10757668 in exon 2 splice acceptor (*premixed with exon1B-exon2 probe below*) | AN7DZHR | Thermo | 4332077 | FAM/VIC |
| 1B Allele Specific | Exon 1B-exon 2 junction (NM_018325.4) targeting SNP rs10757668 in exon 2 splice acceptor (*premixed with exon1A-exon2 probe above*) | AN9HU3N | Thermo | 4332077 | FAM/VIC |
| UBE2D2 HEX | Housekeeping gene | UBE2D2, Human (HEX); dHsaCPE5036421 | Biorad | 10031255 | HEX |
| UBE2D2 FAM | Housekeeping gene | UBE2D2, Human (FAM) | Biorad | 10031252 | FAM |

**Extended Data Figure 13.**
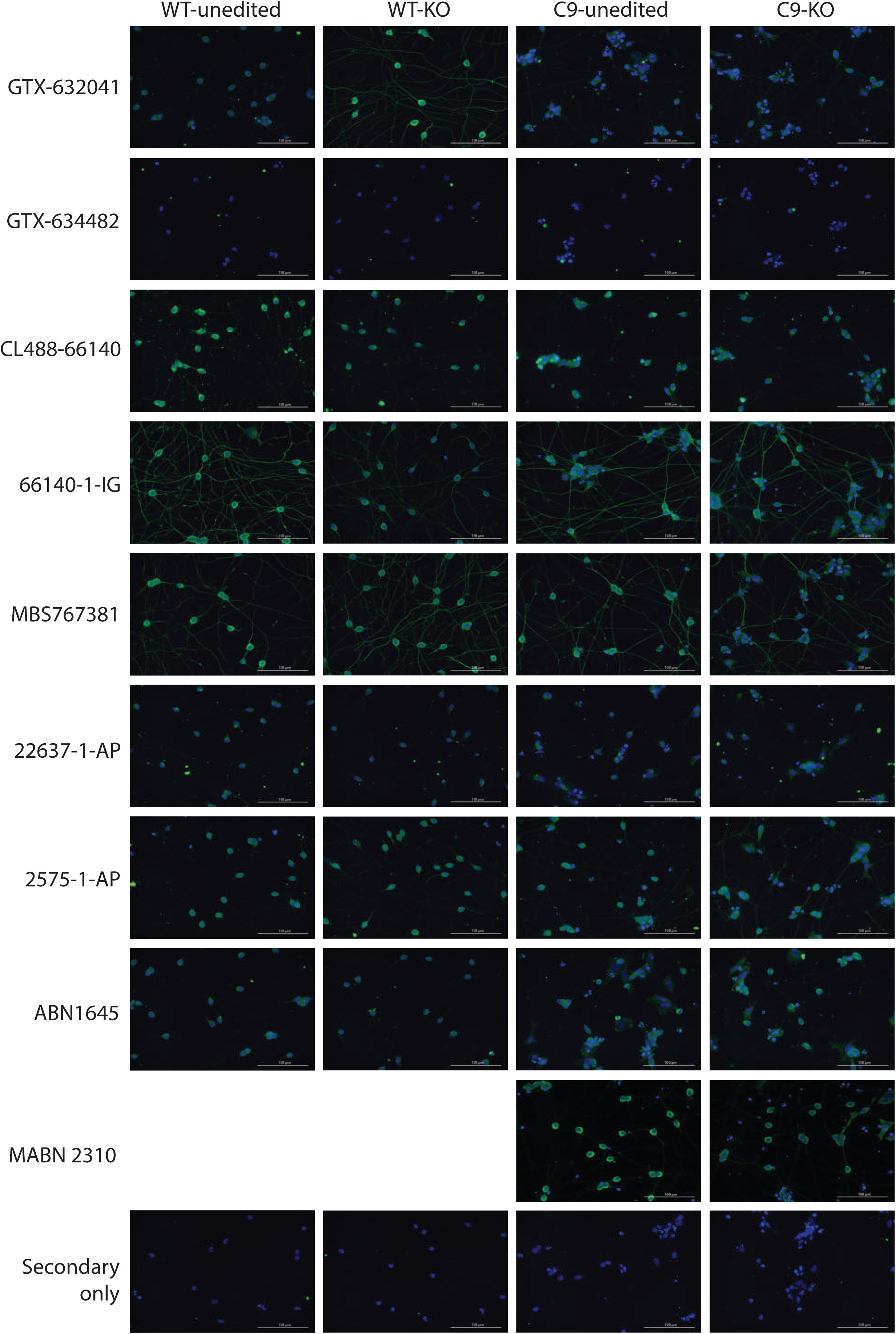
Nine commercially available C9orf72 antibodies are not specific for C9orf72 in iPSC-derived neurons by immunocytochemistry. (A) We found that commercially available C9orf72 antibodies were not specific for C9orf72 by comparing staining patterns in knock-out lines (WT-KO and C9-KO) to unedited cells (WT-unedited and C9-unedited). Blue = DAPI. Green = staining from antibodies listed in Supplementary Table 4. Scale bar = 100uM.

**Supplementary Table 4.** Commercially available C9orf72 antibodies tested in Extended Data Fig. 13, with the concentrations used.

| Primary Antibody | Company | Primary Antibody Dilution | Secondary Antibody | Secondary Antibody Dilution |
| --- | --- | --- | --- | --- |
| GTX-632041 | GeneTex | 1:100 | Goat anti-mouse IgG (H+L)<br>Alexa Fluor 488 | 1:500 |
| GTX-634482 | GeneTex | 1:100 | Goat anti-mouse IgG (H+L)<br>Alexa Fluor 488 | 1:500 |
| CL488-66140 | Proteintech | 1:50 | Goat anti-mouse IgG (H+L)<br>Alexa Fluor 488 | 1:500 |
| 66140-1-IG | Proteintech | 1:500 | Goat anti-mouse IgG (H+L)<br>Alexa Fluor 488 | 1:500 |
| MBS767381 | MyBioSource | 1:50 | Goat anti-mouse IgG (H+L)<br>Alexa Fluor 488 | 1:500 |
| 22637-1-AP | Proteintech | 1:500 | Goat anti-rabbit IgG (H+L)<br>Alexa Fluor 488 | 1:500 |
| 25757-1-AP | Proteintech | 1:300 | Goat anti-rabbit IgG (H+L)<br>Alexa Fluor 488 | 1:500 |
| ABN1645 | Sigma-Aldrich | 1:500 | Goat anti-rabbit IgG (H+L)<br>Alexa Fluor 488 | 1:500 |
| MABN2310 | Sigma-Aldrich | 1:500 | Goat anti-rabbit IgG (H+L)<br>Alexa Fluor 488 | 1:500 |

**Extended Data Figure 14.**
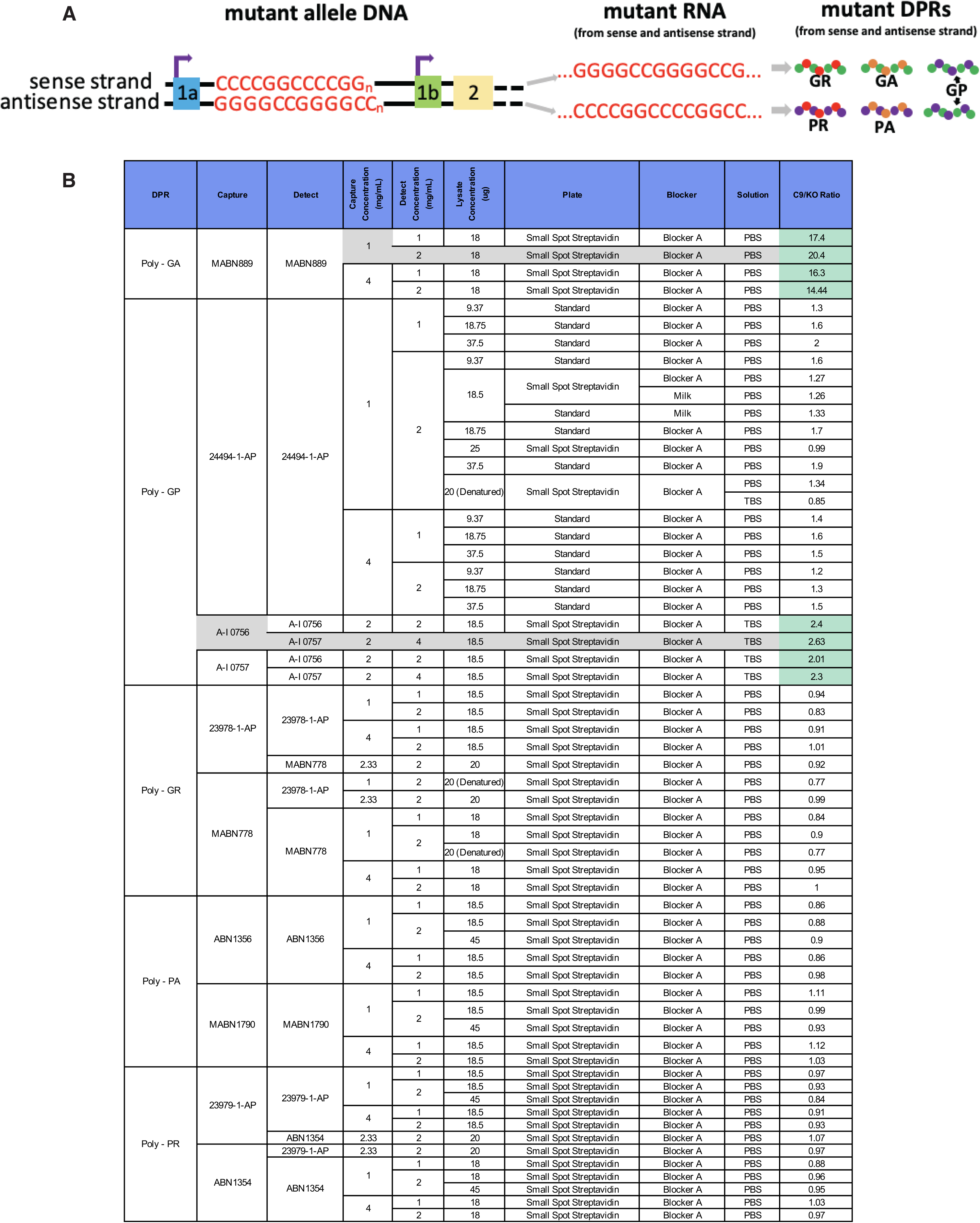
Two of 10 antibodies tested were specific for dipeptide repeats (DPRs) from the C9orf72 mutant line compared to baseline noise as determined by KO control. (A) Schematic of expression of sense and antisense repeat expansion in RNA and through non-canonical repeat-associated non-AUG (RAN) translation to form mutant dipeptide proteins. 5 total DPRs are formed: poly-GR and poly-GA from the sense strand, poly-PR and poly-PA from the antisense strand and poly-GP from both the sense and antisense strands. (B) We tested 10 antibodies raised against DPRs in varying combinations and concentrations using sandwich ELISA on the MSD platform. Concentrations of capture and detect antibodies and lysate concentration from 2-week old neurons are noted. We compared our C9-unedited line harboring an expanded repeat to C9-KO. The 21-21 kb KO stretched from 5’ to the *C9orf72* gene through the transcriptional start sites and the repeat region and included exons 1-3. Antibody combinations that generated a ratio of signal from the C9-patient vs KO line greater than 2 (highlighted in green) were used to generate Figure 3 of the main text (the corresponding conditions used are highlighted in grey). Most antibodies generated noise that was similar between KO and C9-patient lines (ratio ∼1).

## Online methods

### Cell line generation, maintenance and determination of editing efficiencies

We used iPSC generated by others^42, 61, 62^ from patients harboring the C9orf72 mutation and a control cell line without mutation^81^ (WT-control). We maintained iPSCs in mTesR plus plated on Matrigel (Corning 356231), passaging at 60-80% confluency. All cell lines had a normal karyotype and negative quarterly mycloplasma testing.

We first knocked-in the inducible motor neuron transcription factor transgene cassette^51, 82^ in the CLYBL safe-harbor locus of a C9-patient line using spCas9 and ATGTTGGAAGGATGAGGAAA gRNA. This transgene includes human NGN2, ISL1, LHX3 (hNIL) under the TET operator and is inducible by doxycycline, mCherry (for positive selection) and neomycin antibiotic resistance (for negative fluorescence). Red-fluorescing cells were sorted via FACS to isolate single, live cells. Each resulting clonal cell line was analyzed for incorporation of the transgene in the CLYBL locus by PCR (left homology arm junction primers CAGACAAGTCAGTAGGGCCA and AGAAGACTTCCTCTGCCCTC) with preservation of one of the alleles (CLYBL wild-type primers TGACTAAACACTGTGCCCCA and AGGCAGGATGAATTGGTGGA). We used Copy Number Variation (CNV) ddPCR to pick a clone with a single transgene insertion of the hNIL plasmid (Nemomycin primers CATGGCTGATGCAATGCG and TCGCTTGGTGGTCGAATG, probe FAM; Primers UBE2D2 – Bio-Rad 10031255, probe HEX) to mitigate the risk of integration of the transgene at genomic loci other than CLYBL.

To engineer each iPSC line we used HiFi spCas9 protein (Macolabs, UC Berkeley) and two gRNAs (**Supplementary Table 1**) to create an excision, using our published protocol^83^. gRNAs were designed to have no exact off-target matches and the lowest predicted off-targets using CRISPOR (Homo sapiens – USCS Dec. 2013 (GRCh38/hg38))^46^. gRNAs were ordered from IDT or Synthego. Cas9-gRNA RNP (spCas9 (40µM), sgRNA (100µM)) was delivered by nucleofection (Lonza AAF-1002B, Lonza AAF-1002X, Pulse Code = DS138) to 350,000 iPSCs suspended in 20 µl of P3 Buffer. The cells were recovered with mTesR plus supplemented with ROCK1 inhibitor (Selleckchem S1049) at 10 µM and Clone R (Stemcell 05888). Approximately 50% of iPSCs died within the first 24 hours of electroporation, as expected. Following a 48-72 hour recovery, we collected the pool of edited cells and either hand-picked 48 clones or sorted single live cells via FACS to a single well on a 96-well plate. Single cell sorting was performed using a BD FACSAria Fusion (Beckton Dickinson) by the Gladstone Flow Cytometry Core. The QC alignment of each laser was verified with Cytometer Setup and Tracking Beads (Becton Dickinson) before sample acquisition. A forward scatter threshold of 15,000 was set to eliminate debris from list mode data, and a fixed number of events was collected. In some experiments mCherry fluorescence (excitation 561nm, emission 610nm) was also used to define sorting parameters. Drop delay determination and 96-well plate set-up setup was done using Accudrop beads (Becton Dickinson). Gating on forward scatter area versus height and side scatter area versus height was used to make the single cell determination. The specifications of the sort layout included single cell precision, 96-well collection device and target event of 1. After cultures reached 60-70% confluency, each well was split into two wells of a new 48- or 96-well plate, one for sequencing and the other to continue the cell line. We screened clones based on the presence of an excision band using PCR (primers and expected band size from **Supplementary table 1**). We also performed PCR across each the 5’ and 3’ cut site (**Supplementary Table 1**), with one primer site located inside the excision region, to ensure absence of a band (for homozygous edits) or presence of the WT allele (for heterozygous edits). For all lines except C9-REx, we then Sanger sequenced the excision band (MCLAB). If the sequence was ambiguous (i.e., had overlapping nucleotide reads at the same mapped nucleotide position) we subcloned the line to achieve clone purity and clean sequencing. All lines were karyotyped (WiCell or Cell Line Genetics) after editing.

For all lines except C9-REx, editing efficiency was determined based on the PCR amplification of an excision band, and in the case of homozygous excisions, the absence of the WT band, in each clone (48 hand-picked clones or 96 single-cell FACS sorted clones). See **Supplementary Table 1** for primers. For C9-REx we could not use this approach since PCR could not amplify the large repeat expansion, and hence could not distinguish clones with excision of both the mutant and WT allele from clones with excision of the WT allele only. Therefore, we used single-molecule sequencing of clonal REx lines to determine the percentage of clones with an excision of the repeat expansion region (as described below).

### PacBio single molecule sequencing to size the repeat expansion and detect repeat expansion excision (C9-REx)

Because polymerase amplification fails to accurately size the entirety of the *C9orf72* GC rich repetitive region, we use single molecule sequencing^58, 84^ of a genomic region containing the repeat region. We collected high molecular weight DNA using Genomic Tip (Qiagen 10243) and confirmed absence of smearing by running the DNA on a 1.5% agarose gel. The Gladstone Genomics Core performed library preparation according to the “No Amp Targeted Sequencing” published protocol^58, 84, 85^ using 3-5 µg of DNA per sample as measured by Qubit. Briefly, we blocked the free ends of purified genomic DNA and then excised the gene region of interest using spCas9, a gRNA targeting 5’ to the repeat expansion (GGAAGAAAGAATTGCAATTA) and a gRNA targeting 3’ to the repeat expansion (TTGGTATTTAGAAAGGTGGT). Excising the genomic region harboring the repeat expansion yields a 3639 bp fragment from the WT allele and a fragment of variable size from the mutant allele depending on the size of the CCCCGG repeat. We then ligated adapters and barcodes to blunt free ends and sequenced 3-5 barcoded lines per SMRT Cell on either a Sequel I or Sequel II sequencer. We used a 3-pass filter such that each molecule of DNA had to be sequenced at least 3 times to be included in analysis. We compared repeat counts from sequencing to Southern blot, performed by Celplor using 20 µg of input DNA and the previously published protocol^57^.

### iPSC differentiation into motor neurons

We used the hNIL transgene cassette TET-on system in the CLYBL safe-harbor locus of a C9-patient line and WT-control line. Introduction of doxycycline for 3 days induced the expression of 3 human transcription factors: NGN2, ISL1, LHX3. We followed the previously published protocol^51, 82^ with notable exceptions, including higher concentrations of the growth factors BDNF, GDNF and NT-3 (each at 20 ng/ml). Our detailed protocol is published^86^.

### RNA quantification by ddPCR

2-week old induced neurons were lysed with papain (Worthington LK003178) and RNA was isolated using Quick-RNA Microprep Kit (Zymo R1051). cDNA was synthesized using iScript™ Reverse Transcription Supermix (Biorad 1708841) and 500 ng of RNA. ddPCR was run with 3 technical replicates of each of 3 biologic replicates (independent wells of differentiated neurons) on the QX100 Droplet Reader (Bio-Rad 186-3002). Each ddPCR reaction consisted of 12.5 uL of 2x SuperMix for Probes (no dUTP) (Bio-Rad 186-3024), primer/probe (see **Supplementary Table 3**), 5 ng of cDNA, and nuclease-free water up to 25 µL. Droplets were generated with QX 100 Droplet Generator (Bio-Rad 186-3001) and 20 µL of the reaction mixture with 70 µL of oil. The ddPCR reactions were run in a Deep Well C1000 Thermal Cycler (Bio-Rad 1851197) with the following cycling protocol: (1) 95°C for 10 min; (2) 94°C for 30 s; (3) 58°C for 1 min; (4) steps 2; and 3 repeat 39 times; (5) 98°C for 10 min; (6) hold at 4°C. We thresholded positive samples as those with >10 positive droplets to avoid error due to noise. We quantified positive droplets for each target and normalized the amount to our loading control (*UBE2D2*) (Bio-Rad QuantaSoft™ Analysis Pro Software). We chose this housekeeping gene because its expression level remained stable across iPSCs and differentiated neurons^87^.

For allele-specific expression of exon 1A- and 1B-containing transcripts, we utilizing a coding SNP in the exon 2 splice acceptor (rs10757668) in our patient line. We centered our ddPCR probe on this SNP and used the same primers as above to amplify the exon 1A-exon 2 (Thermo, 4332077) and exon 1B-exon 2 junctions (Thermo, 4332077) (**Supplementary Table 3**). Expression from each allele was quantified in a single reaction and reported as a ratio.

### C9orf72 protein quantification by Simple Western

We performed protein quantification by streptavidin-based Simple Western^88^ capillary reaction (WES; Bio-Techne) according the manufacturers protocol (Jess & Wes Separation Module SM1001 to SM1012^89^), with the following specifications: protein was collected from cultured neurons 2 weeks post-induction in RIPA buffer with protease inhibitor and sonicated for 5 min, and denatured at 90°C for 10 min. 0.3 µg/µl protein from each sample was mixed with 1 µl 5x Master Mix and 0.1x Sample Buffer (EZ Standard Pack PS-ST01EZ-8) to a total volume of 5 µl. 3 µl of this mix was loaded per sample onto a 12-230 kDa plate (ProteinSimple SM-W004-1). Primary antibodies were mouse anti-C9orf72 (GeneTex, GTX634482) at a 1:100 dilution and rabbit anti-GAPDH (AbCam, AB9485) at 1:1000 dilution (total volume 10 µl per lane). Duplexed secondaries included 9.5 µl of mouse (ProteinSimple, DM-002) and 0.5 µl of 20x anti-rabbit (ProteinSimple, 043-426) per lane. Reaction times: 25 min separation time at 375 V, 5 min antibody dilutant time, 30 min primary antibody, 30 min secondary antibody; quantification at 4 seconds of detection (high dynamic range). Under these optimized conditions, each antibody produced a single peak at 57 kDa (C9orf72) and 42 kDa (GAPDH). Area under the curve was quantified for each peak and C9orf72 AUC was normalized to GAPDH AUC for each sample. Averages across 3 biological replicates (independent wells of neuronal differentiation) of neurons from each edited cell line aged 14 days post-induction were compared to the average protein expression of their respective unedited controls.

### Dipeptide repeat quantification by Meso Scale Discovery (MDS) sandwich ELISA

We found 2 antibody combinations to be specific for detecting DPRs in 14-day-old iPSC-derived neurons harboring the *C9orf72* repeat expansion (**Extended Data Fig. 14**). We followed the manufacturer’s protocol for the Small Spot Streptavidin Plate (L45SA, MSD). Poly-GA was detected using anti-GA antibody (MABN889, Millipore) at 1 mg/ml (capture) and 2 mg/ml (detect) final concentration and 18 µg total protein per sample (blocking buffer A, solution PBS). Poly-GP was detected using anti-GP antibody (affinity purified TALS828.179 from TargetALS, purification lot A-I 0757 and stock concentration 1.39 mg/ml). A-I 0757 anti-GP antibody was used at a final concentration of 2 mg/ml capture and 4 mg/ml detect with 18.5 µg total protein per sample (blocking buffer A, solution TBS). The plate was coated with capture antibody overnight at 4°C with no agitation. The plate was blocked with 3% MSD Blocker A (R93BA, MSD) in 1X DPBS for 1 hour at 750 rpm, then incubated for 1 hour with protein lysate at 750 rpm at room temperature. Detection antibody was added after the lysate for 1 hour. Washes were performed between steps thrice with 1X DPBS + 0.05% Tween-20. MSD Read Buffer A (R92TG, MSD) was added to the plate before being immediately placed in the MSD Model 1250 Sector Imager 2400 plate reader. Signal was calculated by comparing luminescence intensity for each control or edited patient line to background (*i.e.,* C9-KO line), data was presented as a fold change above C9-KO baseline/background level. Our detailed MSD sandwich ELISA protocol is published^90^.

### TDP-43 immunocytochemistry and quantification

7-week-old neurons were fixed by adding 4% PFA directly to culture media for 30 min followed by 3 PBS washes of 10 min each. Cells were permeabilized by 1X DPBS 0.1% Triton-X in 3 washes of 10 min each at room temperature and blocked with 1X DPBS 0.1% Triton-X + 5% BSA for 1 hour at room temperature. Primary antibodies: rabbit anti-TDP43 (10782-2-AP, Proteintech) at 1:500, beta-III-tubulin (480011, Invitrogen) at 1:250. Primary antibodies were incubated overnight at 4°C. Secondary antibodies included Goat anti-rabbit Alexa Fluor 488 nm and Goat anti-mouse Alexa Fluor 594 nm. Secondary antibodies were incubated at room temperature for 1 hour. DAPI (D1306, ThermoFisher Scientific) was added to the penultimate of five, 5 min PBS washes. Our detailed immunocytochemistry protocol is published^91^. After staining, cells were scanned on the ImageXpress Micro Confocal (Molecular Devices). TDP-43 cells were quantified using Elements AI (Imaging Software NIS-Elements AR 5.30.04 64-bit). We trained the software to differentiate TDP-43-positive cells with or without nuclear TDP-43 on images from independent differentiations not included in the quantification. A blinded observer hand-classified and hand-counted TDP-43-positive cells and confirmed the trends detected by AI.

## Acknowledgments

C.D.C. Is supported by NIH K08-NS112330KO8, UCSF CTSI TL1 Fellowship 5TL1TR001871-04, Bright Focus Foundation A20201490F, Weill Institute for Neurosciences and UCSF Memory & Aging Center. C.D.C. acknowledges a gift from the Ludwig Family Foundation. C.D.C and B.R.C. are supported by RF1-AG072052. B.R.C. is supported by (R01-HL130533, R01-HL13535801, P01-HL146366). B.R.C. acknowledges support through a gift from the Roddenberry Foundation, Keenon Werling and Pauline and Thomas Tusher. We thank F. Chanut, L. Judge, H. Watry, J. Perez-Bermejo, Z. Nevin, H. Sun and M. Bernardi for technical advice and editorial comments.

## Author Contributions

C.D.C, B.R.C. conceived of the study. C.D.C designed the studies, analyzed and interpreted the data and wrote the manuscript. K.C.K. created and implemented AlleleAnalyzer. A.S. and K.G. generated the cell lines. A.S., K.G., A.B., H.W. performed and analyzed ddPCR. K.G., M.S. performed and analyzed DPR MSD. TDP-43 staining and quantification was performed by M.S. A.B. performed C9orf72 ICC. Y-C.T. and J.Z. performed and analyzed single-molecule sequencing.

